# Pharyngeal pouches provide a niche microenvironment for arch artery progenitor specification

**DOI:** 10.1101/2020.05.07.083493

**Authors:** Aihua Mao, Linwei Li, Jie Liu, Mingming Zhang, Guozhu Ning, Yu Cao, Qiang Wang

## Abstract

The paired pharyngeal arch arteries (PAAs) are transient blood vessels connecting the heart with the dorsal aorta during embryogenesis. Although PAA malformations often occur along with pharyngeal pouch defects, the functional interaction between these adjacent tissues remains largely unclear. Here we report that the ablation of pouches in zebrafish embryos impairs PAA progenitor specification and leads to the absence of PAA structures. Through time-lapse imaging studies, we reveal that the segmentation of pharyngeal pouches coincides spatiotemporally with the emergence of PAA progenitor clusters. These pouches physically associate with pharyngeal mesoderm in discrete regions and provide a niche microenvironment for PAA progenitor commitment by expressing BMP proteins. Specifically, tissue specific knockdown experiments demonstrate that pouch-derived BMP2a and BMP5 are the primary niche cues responsible for activating the BMP/Smad pathway in pharyngeal mesoderm, thereby promoting progenitor specification. In addition, BMP2a and BMP5 play a primary inductive function in the expression of the *cloche* gene *npas4l* in PAA progenitors. Mutation of the *cloche* locus represses the specification of PAA progenitors and generates ectopic muscle precursors in the pharyngeal mesoderm. Therefore, our results support a critical role of pharyngeal pouches in establishing a progenitor niche for PAA morphogenesis via BMP2a/5 expression.

## Introduction

During vertebrate development, the pharyngeal arch arteries (PAAs), also known as aortic arch arteries, are transient embryonic blood vessels that connect the heart to the dorsal aorta and establish the circulatory system (Hiruma et al., 2002). These arteries form in a cranial-to-caudal sequence, followed by the regression of the first and second PAAs, whereas the PAAs 3, 4 and 6 undergo asymmetric remodeling and contribute to the carotid arteries and great vessels of the heart, including the aorta and pulmonary arteries (Congdon, 1922; Hiruma et al., 2002). Improper embryonic development of the PAAs may cause life-threating congenital cardiovascular defects that are frequently of unknown etiology (Abrial et al., 2017; Srivastava, 2001). In the past ten years, the regulatory mechanisms involved in PAA remodeling have been studied extensively (Abu-Issa et al., 2002; Kameda, 2009; Liu et al., 2004; Papangeli and Scambler, 2013; Watanabe et al., 2010), however, the cellular events and genetic control of PAA formation are just beginning to be unveiled.

As one of the vertebrate classes, zebrafish embryo exhibits very similar processes of PAA morphogenesis, despite the absence of aortic arch remodeling (Anderson et al., 2008; Isogai et al., 2001). During zebrafish mid-somitogenesis, a common progenitor population, the lateral pharyngeal lineage that gives rise to PAAs, head muscles (HM) and cardiac outflow tract (OFT), segregated from the cardiac precursors in the heart field (Guner-Ataman et al., 2018). These common progenitors are housed in different pharyngeal arches and exhibit distinct gene expression profiles prior to the morphogenesis of PAAs, HM and cardiac OFT (Nagelberg et al., 2015; Paffett-Lugassy et al., 2017). In particular, the common progenitors located in PAAs 3-6 condense into *nkx2.5^+^* clusters in a craniocaudal sequence and then gives rise to PAA endothelium, implying the progressive emergence of PAA progenitors (Paffett-Lugassy et al., 2013). Specifically, cell lineage tracing experiments showed that progenitors for PAAs 5 and 6 were specified from the pharyngeal mesoderm after 30 hpf, a time point far away from the segregation of the common progenitors (Paffett-Lugassy et al., 2013). These observations suggest that these common progenitors might undergo further specification in the pharyngeal region, a hypothesis that remains to be evaluated experimentally.

Transcription factors *etv2* and *scl* have been shown to be essential for the initiation of the angioblast program (Sumanas and Lin, 2006). Interestingly, *nkx2.5*^+^ acts upstream of *etv2* and *scl* to promote the transition from PAA progenitor to angioblast, however, it is dispensable for PAA progenitor specification (Paffett-Lugassy et al., 2013). In addition, chemical inhibition experiments suggest that transforming growth factor β (TGF-β) signaling is required for the differentiation of PAA progenitors into angioblasts (Abrial et al., 2017). However, the molecular mechanism underlying PAA progenitor specification in the pharyngeal region has yet to be fully investigated.

Endodermal pouches are a series of outpocketings budding from the developing foregut (Graham and Smith, 2001). Interestingly, affected arch arteries often occur simultaneously with pouch defects, possibly because of their close physical relation and potential interactions during development (Li et al., 2012; Wendling et al., 2000). Because the pharyngeal pouches express several signaling molecules that participate in the patterning of the pharyngeal skeleton and in the specification of the arch-associated ganglia, their roles in aortic arch morphogenesis have been traditionally considered as secondary (Crump et al., 2004; Holzschuh et al., 2005; Ning et al., 2013). Intriguingly, our recent study indicated an indispensable role of PDGF signaling from pharyngeal pouches in the PAA angioblast proliferation (Mao et al., 2019), but whether pouch endoderm directly functions in PAA progenitor specification remains unknown. In this work, we found that the common progenitors for PAAs, HM and cardiac OFT may be further specified into two vascular progenitor subpopulations within the pharyngeal mesoderm: the *nkx2.5^+^* progenitors that give rise to PAAs and *nkx2.5^-^* progenitors that generate the connective ventral aorta. Moreover, the cloche gene *npas4-like* (*npas4l*) is expressed in PAA progenitors and mutation of the *cloche* locus induce a transformation of the pharyngeal mesoderm from a given endothelial fate to a muscle cell fate, indicating the pharyngeal mesoderm harbor multiple differentiation potential into at least the endothelial and muscular lineages. During PAA morphogenesis, the pouches interact with the pharyngeal mesoderm in a cranial to caudal sequence. Nitroreductase (NTR)-mediated tissue ablation experiments illustrated that the pouches, while not the cranial neural crest cells that give rise to the pharyngeal skeletons, play an essential function in PAA progenitor specification. We further found that pouch-derived BMP signaling is necessary for endothelial lineage commitment within the pharyngeal mesoderm.

## Results

### ZsYellow^+^ pharyngeal mesoderm contains distinct vascular progenitor subpopulations

Previous studies have demonstrated that PAAs originate from a fraction of *nkx2.5*^+^ cells within the heart field through vasculogenesis (Guner-Ataman et al., 2018; Nagelberg et al., 2015; Paffett-Lugassy et al., 2013). To meticulously observe cell behaviors during the formation of these PAAs, time-lapse confocal imaging studies were performed in *Tg(nkx2.5:ZsYellow)* embryos from 22 hpf. At this time point, some cells in the ZsYellow^+^ pharyngeal mesoderm started to pile up in the region corresponding to the ventral root of the prospective third aortic arch, and then sprouted dorsally by 24 hpf (S1A Fig). This process was repeated for PAAs 4-6 in a cranial to caudal sequence from 28 to 42 hpf (S1A Fig), which is consistent with previous observations that PAA progenitors condense into clusters concomitant with pharyngeal segmentation (Paffett-Lugassy et al., 2013).

During somitogenesis, the common progenitors of PAAs, HM and cardiac OFT are specialized and remain lateral with consecutive *nkx2.5* expression when cardiac precursors migrate medially and contribute to the heart (Guner-Ataman et al., 2018; Paffett-Lugassy et al., 2013). Interestingly, the PAA progenitor clusters sequentially emerged at discrete positions in the pharyngeal mesoderm to form aortic arches connecting the lateral dorsal aorta and ventral aorta to support blood flow, whereas the ZsYellow*^+^* cells between the PAA progenitor clusters seemed to preserve their locations and would not contribute to PAAs (S1A and S1B Fig). These observations support that the pharyngeal mesoderm within pharyngeal arches 3-6, where PAA progenitors gives rise to aortic arches, might be further specified into different subpopulations.

To test this hypothesis, we evaluated the expression pattern of *nkx2.5*, the specific marker of PAA progenitors from 28 hpf to 38 hpf (Guner-Ataman et al., 2018; Nagelberg et al., 2015; Paffett-Lugassy et al., 2013). As described in previous reports (Paffett-Lugassy et al., 2013), *ZsYellow* transcripts derived from *nkx2.5* cis-regulatory sequences in *Tg(nkx2.5:ZsYellow*) embryos gradually appeared in the progenitor clusters in a craniocaudal sequence (S1C Fig).

The endogenous *nkx2.5* transcripts in wild-type embryos were also sequentially observed, but eventually decreased when the PAA progenitors differentiated into angioblasts (S1C Fig). Importantly, the transcripts of both *ZsYellow* and *nkx2.5* were segregated in different domains (S1C Fig). To further investigate this observation, we combined immunofluorescence and fluorescence *in situ* hybridization experiments, and found that most of the *nkx2.5^+^* progenitors were restricted to the PAA clusters within ZsYellow^+^ pharyngeal mesoderm, and the *etv2^+^* and *scl^+^* PAA angioblasts located in the ventral root of each sprouts (Fig 1A and 1B). Taken together, these results show that the pharyngeal mesoderm within pharyngeal arches 3-6 is composed of distinct subpopulations with or without *nkx2.5* expression.

**Fig 1.**
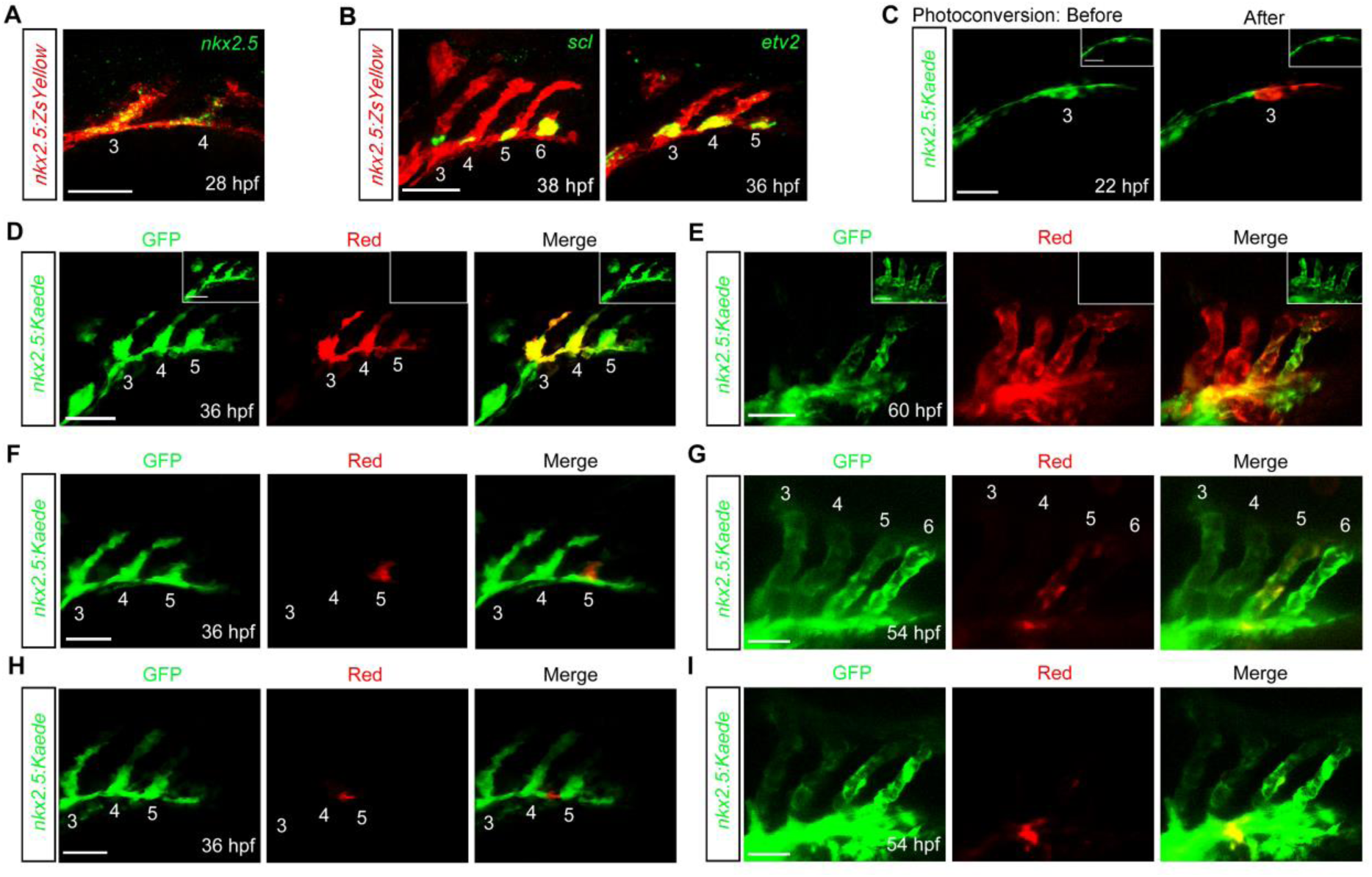
Pharyngeal mesoderm contains two vascular progenitor subpopulations. (A-B) The expression patterns of PAA progenitor marker *nkx2.5* (A) and angioblast markers *scl* and *etv2* (B) in *Tg(nkx2.5:ZsYellow)* embryos were detected by fluorescent *in situ* hybridizations. Embryos were first subjected to fluorescent *in situ* hybridizations and then to immunostaining with anti-ZsYellow antibody. Numbers indicate PAA clusters or sprouts. Lateral views with anterior on the left. Scale bar, 50 μm. (C-E) The Kaede proteins in PAA cluster 3 and the subsequent posterior pharyngeal mesoderm in the right side of the *Tg(nkx2.5:kaede)* embryos were photoconverted at 22 hpf (C). Cells in the left side of the pharyngeal mesoderm remained un-photoconverted as an internal control (inset). Embryos were subsequently imaged in the green and red channels at 36 hpf and 60 hpf (D and E). Numbers indicate PAA clusters or sprouts. Scale bars, 50 μm. (F-I) Localized photoconversion of PAA cluster 5 (F) or Kaede^+^ cells between cluster 4 and 5 (H) at 36 hpf. Red, green and merged images of the same embryos at 54 hpf are shown in (G) and (I). Numbers indicate PAA clusters. Scale bars, 50 μm.

The above findings raised an interesting question as to whether these subpopulations in the pharyngeal mesoderm undergo distinct cell fates. To answer this question, we performed lineage-tracing analysis in *Tg(nkx2.5:kaede)* embryos, where the pharyngeal mesodermal cells expressing photo-convertible Kaede proteins that could instantly switch from green to red fluorescence following ultraviolet light exposure (Guner-Ataman et al., 2013). In the first set of experiments, the Kaede proteins in the PAA progenitor cluster 3 and the subsequent posterior pharyngeal mesoderm located on the right-side of the embryo were photoconverted at 22 hpf, whereas the pharyngeal mesoderm on the left side remained unconverted as an internal control (Fig 1C). As expected, the derivatives of the photoconverted cells were found in the sprouts of PAAs 3-5 at 36 hpf and in the endothelium of the aortic arches 3-6 as well as the ventral aorta at 60 hpf (Fig 1D and 1E). Interestingly, less red fluorescence and more green fluorescence were observed in the cells of caudal PAAs 5-6 and the posterior portion of ventral aorta (Fig 1E), suggesting that the progenitors of these structures are specified at later stages and might undergo more proliferation during vasculogenesis after photoconversion. Nevertheless, these results indicate that the endothelial cells of PAAs and ventral aorta originate from the Kaede^+^ pharyngeal mesoderm.

Next, we specifically photoconverted the Kaede proteins in PAA cluster 5 at 36 hpf (Fig 1F). The photoconversion process resulted in red derivatives in PAA 5 at 54 hpf, but not in other PAAs (Fig 1G). Interestingly, a few cells with red fluorescence were observed in the junction of PAA 5 and ventral aorta (Fig 1G), suggesting the occurrence of endothelial cell rearrangements during blood vessel fusion (Herwig et al., 2011). In contrast, the photoconversion of Kaede^+^ cells located between PAA cluster 4 and 5 led to red derivatives housed specifically in ventral aorta (Fig 1H and 1I). Based on these observations, we concluded that the ZsYellow^+^ pharyngeal mesoderm cells within PAAs 3-6 are specified into two vascular progenitor subpopulations: *nkx2.5^+^* cells that give rise to PAAs and *nkx2.5^-^* cells that generate the connective ventral aorta.

### Pharyngeal pouches have an essential role in PAA progenitor specification

Since the time window of the emergence of ZsYellow^+^ clusters for PAAs is roughly similar to the stages of pharyngeal pouch segmentation, we next aimed to determine the requirement of pouch endoderm during PAA morphogenesis. Firstly, time-lapse recordings of pharyngeal pouch development and PAA formation were performed to more precisely understand these two sets of developmental events in *Tg(nkx2.3:mCherry;nkx2.5:ZsYellow)* embryos, in which pharyngeal pouches were labeled with red fluorescent protein mCherry (Choe et al., 2013). At around 24 hpf, the third pharyngeal pouch appeared to have fully formed and reached the sprouting ZsYellow^+^ cluster that would eventually give rise to PAA3 (Fig 2A). At later stages, the fourth, the fifth and the sixth pouches successively made contact with the developing ZsYellow^+^ clusters for PAAs 4-6 (Fig 2A), indicating a close interaction between the endodermal pouches and the progenitor clusters within pharyngeal mesoderm. Consistent with previous reports (Alexander et al., 1999), depletion of *sox32* in zebrafish embryos by injection antisense morpholinos (MOs) resulted in a lack of early endoderm and endoderm pouches (S2A and S2B Fig). As previously descripted in endoderm-less *bon* mutants (Paffett-Lugassy et al., 2013), the ZsYellow^+^ cells remained in the pharyngeal region in *Tg(nkx2.5:ZsYellow)* embryos injected with *sox32* MO, but the PAAs were completely absent (Fig 2B).

**Fig 2.**
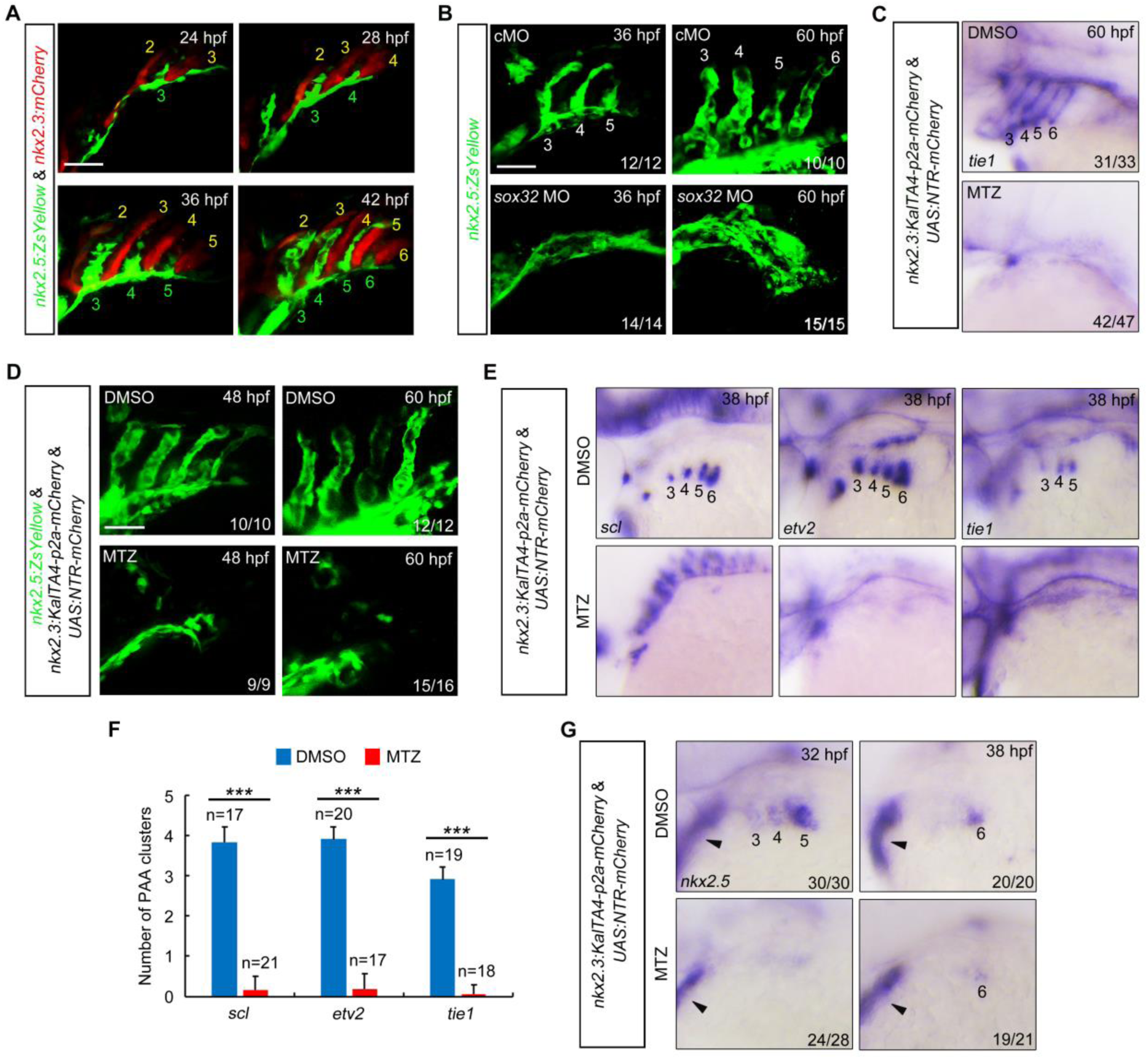
Ablation of pouch endoderm impairs PAA progenitor specification. (A) Confocal images of live *Tg(nkx2.5:ZsYellow;nkx2.3:mCherry)*. The formation of mCherry^+^ pharyngeal pouches (red) coincides with the emergence of ZsYellow^+^ PAA clusters (green). Pharyngeal pouches and PAA clusters or sprouts are numbered with yellow and green Arabic numerals, respectively. Scale bar, 50 μm. (B) Confocal images of live *Tg(nkx2.5:ZsYellow)* embryos show that injection of 8 ng *sox32* MO resulted in complete absence of PAA sprouts at 36 hpf and PAA tubular structures at 60 hpf. The ratios of affected embryos are indicated. Scale bar, 50 μm. (C-D) Embryos were exposed to 10 mM MTZ from bud stage to 38 hpf, and then harvested at the indicated developmental stages for *in situ* hybridization (C) or *in vivo* confocal imaging (D). Numbers in C mark the PAA3-PAA6. Scale bar, 50 μm. (E-F) The *scl*, *etv2* and *tie1* transcripts were evaluated by *in situ* hybridizations in *Tg(nkx2.3:KalTA4-p2a-mCherry;UAS:NTR-mCherry)* embryos treated with DMSO or 10 mM MTZ (E). The average numbers of *scl^+^*, *etv2^+^* and *tie1^+^* PAA angioblast clusters were quantified from three independent experiments and the group values were expressed as mean ± SD (F). Student’s *t*-test, \*\*\**p* < 0.001. (G) Expression of *nkx2.5* in pouch endoderm-depleted embryos. *Tg(nkx2.3:KalTA4-p2a-mCherry;UAS:NTR-mCherry)* embryos were treated with 10 mM MTZ from bud stage to the indicated stages, and then harvested for *in situ* hybridizations. Numbers indicate PAA clusters. Black arrowheads represent the expression of *nkx2.5* in the developing heart.

In order to examine the specific function of pouch endoderm in the establishment of PAAs, we generated a NTR-mediated tissue ablation system, *Tg(nkx2.3:KalTA4-p2a-mCherry;UAS:NTR-mCherry)*, using an optimized Gal4-UAS system to drive NTR protein expression in *nkx2.3*^+^ cells (Curado et al., 2007; Curado et al., 2008; Distel et al., 2009). The *Tg(nkx2.3:KalTA4-p2a-mCherry;UAS:NTR-mCherry)* embryos were exposed to 10 mM MTZ from the bud stage to 36 hpf. Living embryo imaging revealed that over the duration of MTZ treatment, mCherry-expressing pouch endoderm was markedly reduced at 24 hpf and all the pouch structures were successfully ablated at 36 hpf (S3A Fig). Intriguingly, in the pouch endoderm-ablated embryos, the expression of PAA endothelial cell marker *tie1* was absent and the ZsYellow*^+^* pharyngeal mesoderm did not undergo sprouting nor give rise to PAAs (Fig 2C and 2D). Collectively, these results provide strong evidences that pharyngeal pouches are essential for PAA morphogenesis. In addition, MTZ-treated *Tg*(*flk:EGFP; nkx2.3:KalTA4-p2a-mCherry;UAS:NTR-mCherry*) embryos also showed severe defects at these vessels (S3B Fig), indicating the lack of plasticity in the formation of the PAAs when pouch endoderm was ablated (Nagelberg et al., 2015). Furthermore, Cxcl12b, a Cxcr4a ligand derived from the endoderm underlying the lateral dorsal aortae (LDA), has been reported to be required specifically for the formation of LDA (Siekmann et al., 2009). Interestingly, the LDA in pouch endoderm-less embryos displayed no obvious malformations (S3B Fig). This result confirms the specificity of NTR-mediated pouch endoderm ablation in our related experiments.

To determine whether pharyngeal pouches function in PAA progenitor specification, the expression of earlier angioblast lineage markers *scl* and *etv2* and more mature angioblast marker *tie1* was firstly examined in MTZ-treated *Tg(nkx2.3:KaTAa4-p2a-mCherry;UAS:NTR-mCherry)* embryos at 38 hpf. We found that the formation of these angioblast clusters was abolished in embryos treated with MTZ (Fig 2E and 2F). To eliminate the secondary impacts of pharyngeal patterning, we generated a transgenic line, *Tg(sox10:KalTA4-p2a-mCherry)*, in which the red fluorescence of mCherry proteins was expressed in the neural crest cells. We then crossed fishes to generate *Tg(sox10:KalTA4-p2a-mCherry;UAS:NTR-mCherry)* embryos. MTZ treatment from the bud stage induced obvious cell death in the pharyngeal neural crest cells at 36 hpf (S3C Fig). However, the PAA angioblasts developed normally (S3D Fig), implying that pharyngeal neural crest cells are not necessary for the specification and angioblast differentiation of PAA progenitors. We then analyzed the expression of the PAA progenitor marker *nkx2.5* at 32 and 38 hpf and found a significant reduction of *nkx2.5*-expressing progenitors of PAAs 3-6 in the MTZ-treated *Tg(nkx2.3:KalTA4-p2a-mCherry;UAS:NTR-mCherry)* embryos (Fig 2G). Moreover, at 18 hpf, pouch-depletion did not disrupt the segregation of the *nkx2.5^+^* common progenitors that gives rise to PAAs, HM and cardiac OFT (S4A and S4B Fig) (Guner-Ataman et al., 2018). Taken together, these results suggest that pouches are essential for PAA progenitor specification in the pharyngeal region.

### Pharyngeal pouches provide a niche for BMP signal activation in presumptive PAA progenitors

Several genes encoding BMP ligands, including *bmp2a*, *bmp2b*, *bmp4* and *bmp5*, are expressed in the pouch endoderm during pharyngeal segmentation (Holzschuh et al., 2005). Indeed, when pouch endoderm was ablated in *Tg(nkx2.3:KalTA4-p2a-mCherry;UAS:NTR-mCherry)* embryos, expressions of these *bmp* genes were eliminated in the pharyngeal region (Fig 3A). We then examined their potential roles in BMP signal activation in ZsYellow^+^ pharyngeal mesoderm. Immunostaining experiments revealed robust signals of phosphorylated Smad1/5/8, the intracellular effectors for BMP signaling, in the forming ZsYellow^+^ clusters and neighboring cranial neural crest cells of *Tg(nkx2.5:ZsYellow)* embryos (Fig 3B). Additionally, we crossed *Tg(nkx2.5:ZsYellow)* with *Tg(BRE:EGFP)*, a BMP signal activity reporter line (Laux et al., 2011). We observed strong GFP protein expression in the ZsYellow^+^ clusters and the other nearby tissues (Fig 3C). These results demonstrate that BMP/Smad signal is highly activated in the presumptive PAA progenitors.

**Fig 3.**
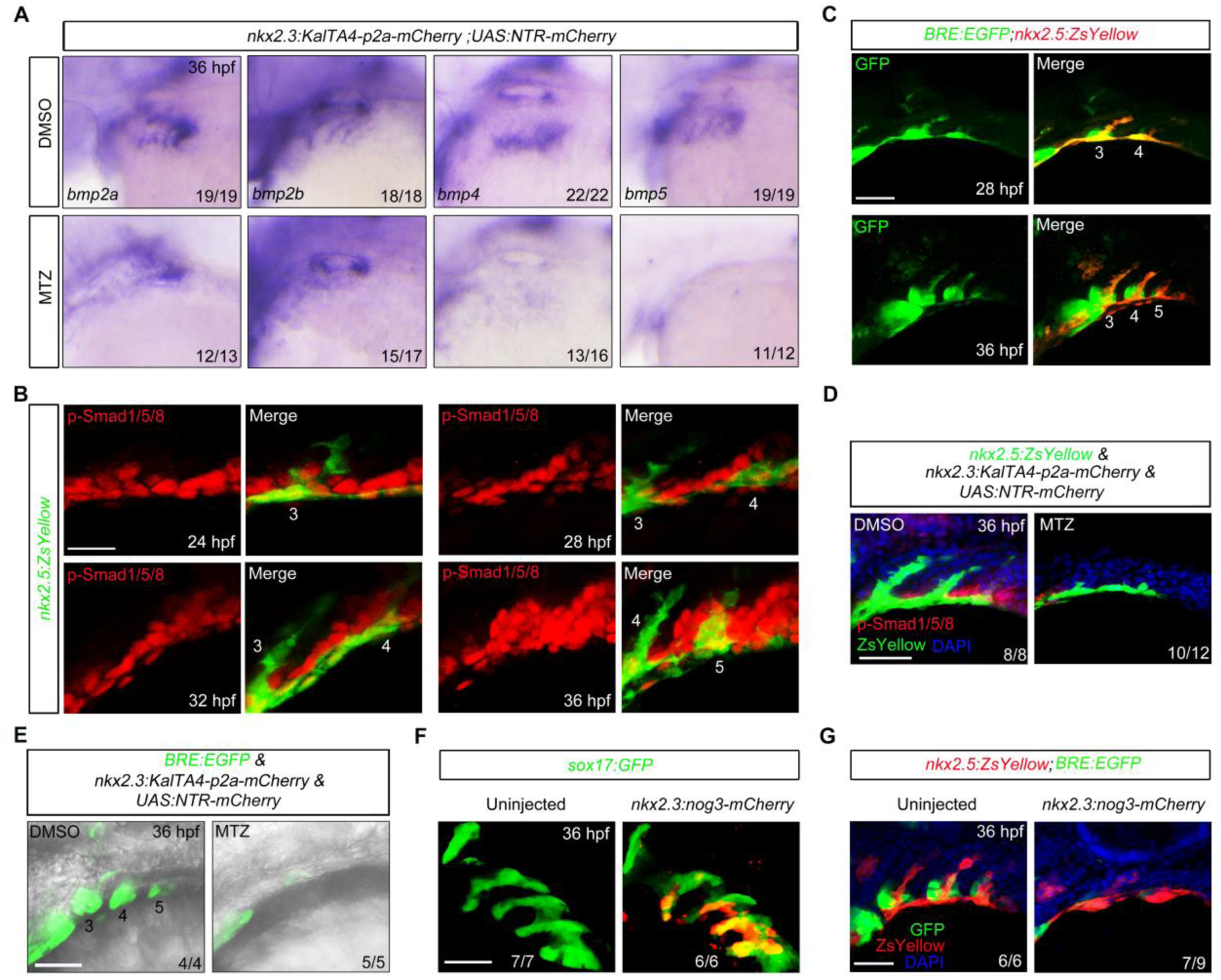
Pouch endoderm is necessary for BMP signal activation in the presumptive PAA progenitors. (A) The *bmp2a*, *bmp2b*, *bmp4* and *bmp5* transcripts were evaluated by *in situ* hybridizations in *Tg*(*nkx2.3:KalTA4-p2a-mCherry*;*UAS:NTR-mCherry)* embryos treated with DMSO or 10 mM MTZ. The depletion of pharyngeal pouches resulted in a significantly reduced expression of *bmp2a*, *bmp2b*, *bmp4* and *bmp5*. (B) BMP signal was dynamically activated in the forming PAA clusters. *Tg(nkx2.5:ZsYellow)* embryos were harvested at indicated stages and subjected to immunostaining for p-Smad1/5/8 (red) and ZsYellow (green). All the embryos were shown in lateral views with anterior on the left. Scale bar, 20 μm. (C)*Tg(BRE:EGFP;nkx2.5:ZsYellow)* embryos at 28 and 36 hpf were immunostained for GFP (green) and ZsYellow (red) to visualize BMP-responsive cells and PAA clusters. Scale bar, 50 μm. (D) p-Smad1/5/8 levels were greatly decreased in pouch endoderm-depleted embryos. *Tg(nkx2.5:ZsYellow;nkx2.3:KalTA4-p2a-mCherry;UAS:NTR-mCherry)* embryos were treated with DMSO or 10 mM MTZ from bud stage to 36 hpf, and then stained for p-Smad1/5/8 (red) and ZsYellow (green). Scale bar, 50 μm. (E) Live confocal images of *Tg(BRE:EGFP;nkx2.3:KalTA4-p2a-mCherry;UAS:NTR-mCherry)* embryos treated with DMSO or 10 mM MTZ from bud stage to 36 hpf. Scale bar, 50 μm. (F) *Tg(sox17:GFP)* embryos were injected with 40 pg *nkx2.3:noggin3-mCherry* plasmids and 200 pg Tol2 transposase mRNA at the one-cell stage, and then harvested for *in vivo* confocal imaging in the green and red channels to visualize pharyngeal pouches (green) and the expression of Noggin3-mCherry fusion proteins. Scale bar, 50 μm. (G) Pouch-derived Noggin3 significantly decreased the BMP signal activity in the pharyngeal region. *Tg(nkx2.5:ZsYellow;BRE:EGFP)* embryos were injected with 40 pg *nkx2.3:noggin3-mCherry* plasmids and 200 pg Tol2 transposase mRNA at the one-cell stage, and then embryos with abundant mCherry fluorescence in the pouches were selected at 36 hpf for immunostaining with anti-GFP and anti-ZsYellow antibodies. Scale bar, 50 μm.

We then ablated the pharyngeal pouches in *Tg(nkx2.5:ZsYellow)* and *Tg(BRE:EGFP)* embryos, respectively. Interestingly, the pouch-depletion led to an evident decrease in BMP signal activity in both ZsYellow^+^ clusters and other pharyngeal tissues (Fig 3D and 3E). These findings imply that pharyngeal pouches function as a niche for activating BMP signals in presumptive PAA progenitors. To further confirm this assumption, *nkx2.3:noggin3-mCherry* plasmids expressing a secreted BMP inhibitory protein Noggin3 (Ning et al., 2013), along with Tol2 transposase mRNA, were co-injected into *Tg(sox17:GFP)* embryos at the one-cell stage. A portion of the injected embryos exhibited uneven, but abundant, mCherry fluorescence in the pouches at 36 hpf (Fig 3F). The same injections were then performed in *Tg(nkx2.5:ZsYellow;BRE:EGFP)* embryos. The embryos with strong mCherry fluorescence in the pharyngeal region were selected at 36 hpf for immunostaining analysis. The pouch-derived Noggin3 significantly inhibited the BMP signal reporter expression (Fig 3G), indicating that the secreted BMP ligands from pharyngeal pouches are the biochemical niche cues that trigger signal activation in presumptive PAA progenitors.

### BMP signaling is required for PAA progenitor specification

To determine whether BMP signaling is required for PAA progenitor specification, we first examined the expression of angioblast marker genes *scl* and *etv2* in embryos treated with 10 μM DMH1, a selective chemical inhibitor of the BMP pathway (Hao et al., 2010; Hao et al., 2014). Interestingly, embryos treated with DMH1 from 20 hpf exhibited impaired angioblast formation in PAA clusters 4-6, whereas the angioblasts in cluster 3 developed normally (Fig 4A and 4B). To further elucidate the role of the BMP pathway in angioblast formation, we treated embryos with DMH1 from 16 hpf, when the common progenitors of PAAs, HM and cardiac OFT have been specified in the ALPM (Guner-Ataman et al., 2018). Noticeably disturbed generation of angioblast clusters 3 and 4-6 was observed (S5A Fig), indicating that BMP signaling is necessary for PAA angioblast formation. However, blocking BMP signaling from such early stages (16 and 20 hpf) led to different severities of pouch defects (S5B Fig), which contributed to the difficulties in distinguishing a direct role of BMP pathway in PAA development. Fortunately, we found that embryos treated with DMH1 from 24 hpf, preceding the formation of PAA cluster 4, showed normal pouch structures (S5B Fig). However, the expression of *scl* and *etv2* was decreased in cluster 4 and completely abolished in clusters 5 and 6 (Fig 4C and 4D). Hereafter, 10 μM Dorsomorphin, another small chemical inhibitor of BMP signaling (Yui et al., 2008), was applied to wild-type embryos from 24 hpf and resulted in similar angioblast phenotypes (Fig 4E and 4F). Consistent with these findings, blocking BMP signaling from 24 hpf greatly reduced the expression of endothelial cell marker *tie1* in the caudal PAAs at 60 hpf (Fig 4G). Similar angioblast and PAA defects were observed in *Tg(hsp70l:dnBmpr1a-GFP)* embryos that were heat-shocked at 24 hpf (S6A-S6C Fig), excluding the potential off-target effects of the pharmacological treatments.

**Fig 4.**
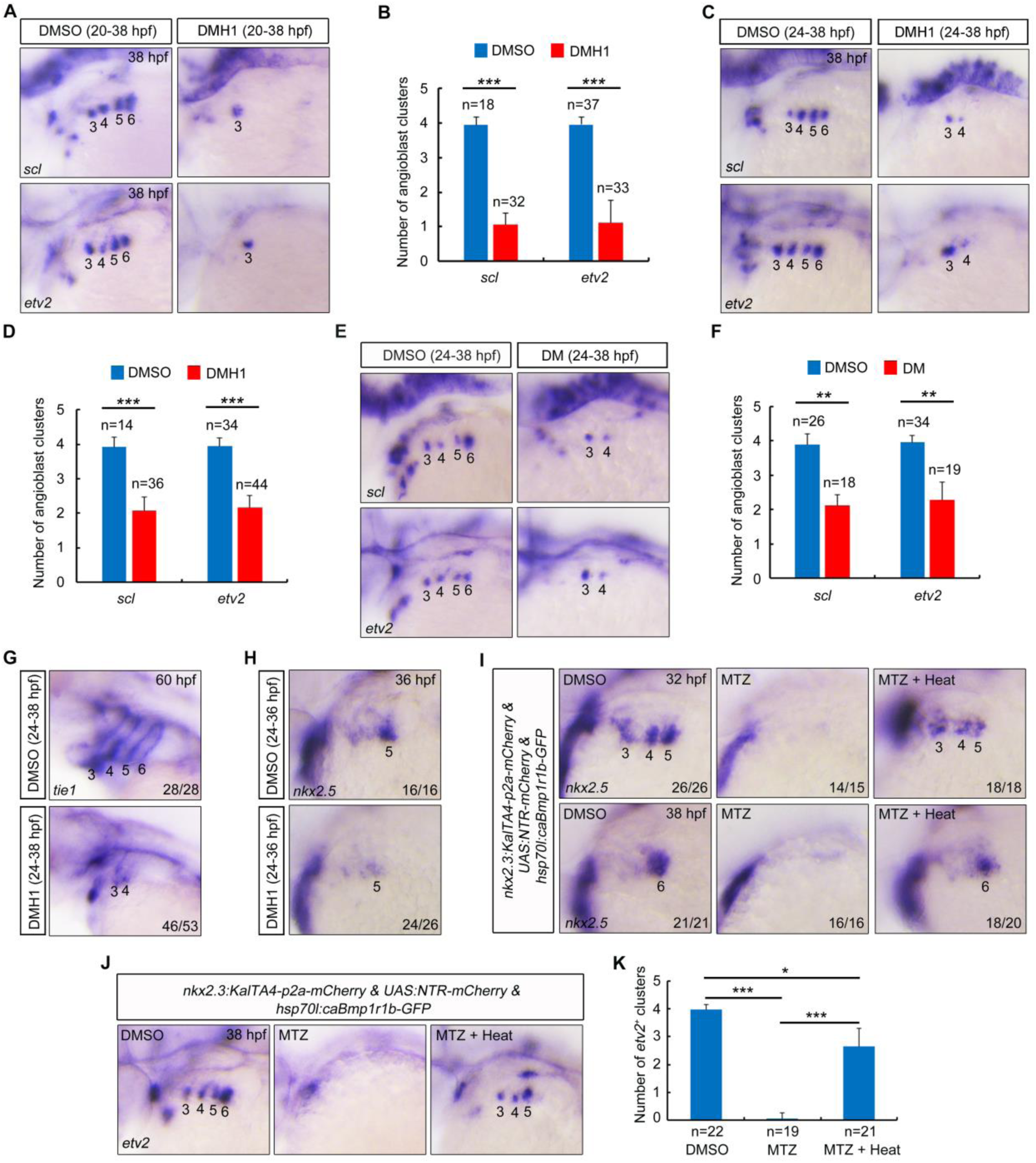
Inhibition of BMP signaling impairs PAA progenitor specification. (A-D) Transcripts of *scl* and *etv2* were evaluated by *in situ* hybridizations in DMSO or DMH1 treated embryos. Embryos were treated with DMSO or 10 μM DMH1 from 20 to 38 hpf (A), or from 24 to 38 hpf (C). Numbers indicate PAA angioblast clusters. The average numbers of *scl^+^* and *etv2^+^* PAA angioblast clusters were quantified in (B) or (D) based on three independent experiments and the group values were expressed as mean ± SD. Student’s *t*-test, \*\*\**p* < 0.001. (E-F)The *scl* and *etv2* transcripts were evaluated by *in situ* hybridization in DMSO or Dorsomorphin treated embryos at 38 hpf. The numbers indicate the PAA angioblast clusters (E). The average numbers of *scl^+^* and *etv2^+^* PAA angioblast clusters were quantified from three independent experiments and the group values were expressed as mean ± SD (F). Student’s *t*-test, \*\**p* < 0.01. (G-H) Wild-type embryos were treated with DMSO or 10 μM developmental stages for *in situ* hybridizations. The numbers indicate the PAAs (G) or PAA progenitor clusters (H). (I-K) Ectopic induction of caBmpr1b rescues the defects of PAA progenitor specification and angioblast differentiation in the pouch-depleted embryos. *Tg(nkx2.3:KalTA4-p2a-mCherry;UAS:NTR-mCherry;hsp70l:caBmpr1b-GFP)* embryos were treated with 10 mM MTZ from bud stage, and heat shocked at 24 hpf for 20 min, and then harvested at the indicated stages for *in situ* hybridizations with *nkx2.5* (I) and *etv2* (J) probes. The average numbers of *etv2^+^* PAA angioblast clusters were quantified from three independent experiments (K). Significance of differences compared with MTZ treatment group was analyzed with Student’s *t*-test, \**p* < 0.05; \*\*\**p* < 0.001.

Next, we analyzed the expression of PAA progenitor marker *nkx2.5* in BMP signal-suppressed embryos at 36 hpf. At this stage, *nkx2.5* transcripts were clearly reduced in PAA clusters 3 and 4 as angioblast differentiated, while they were vigorously expressed in cluster 5 of control embryos (Fig 4H). In contrast, BMP signal inhibition from 24 hpf significantly repressed the *nkx2.5* expression in progenitors residing in cluster 5, indicating a serious imperfection of PAA progenitor specification. Unexpectedly, when *Tg(hsp70l:caBmpr1b-GFP)* embryos were heat-shocked at 24 hpf for 20 min to induce the expression of constitutively active BMP receptor 1b (caBmpr1b), the phosphorylation of Smad1/5/8 was evidently elevated as revealed by immunofluorescence, while PAA progenitor specification and angioblast differentiation remained unchanged (S6D-S6G Fig). These results suggest that BMP signal activation is necessary, but not sufficient, for PAA progenitor specification. Furthermore, the deficiencies in PAA progenitor specification and subsequent angioblast differentiation that induced by pouch-depletion were recovered in heat-shocked *Tg(hsp70l:caBmpr1b-GFP)* embryos (Fig 4I-4K), revealing that specification of PAA progenitors primarily by pouches through activation of BMP signaling in the pharyngeal mesoderm.

### BMP signaling is dispensable for angioblast differentiation, dorsal migration, endothelial maturation and lumen formation during PAA morphogenesis

To explore whether BMP signaling has a role in angioblast differentiation, DMH1 treatment was performed in *Tg(nkx2.5:ZsYellow;gata1:DsRed)* embryos between 30 and 60 hpf, a time window after the specification of progenitors for PAAs 3 and 4. Such DMH1 treatment abolished the formation of PAAs 5 and 6 and led to a lack of blood flow in these caudal PAAs, but had no obvious impact on PAA 3 and 4 (Fig 5A). When the DMH1 treatment was carried out from 38 hpf, a time point when most of the *nkx2.5*^+^ progenitors had accomplished angioblast transition, no obvious defects in PAA development were observed in the resulting embryos (Fig 5A). These results indicate that BMP signaling is crucial for progenitor specification, while dispensable for angioblast differentiation, dorsal migration, endothelial maturation and lumen formation during PAA morphogenesis. It was interesting that if the DMH1 treatment was performed from 30 hpf and then terminated eight hours later, *nkx2.5*^+^ progenitors for the caudal PAAs reappeared at 48 hpf and went on to develop into growing sprouts at 60 hpf (Fig 5A and 5B). Therefore, when BMP inhibition is removed, the pharyngeal mesoderm cells may recover their endothelial potential.

**Fig 5.**
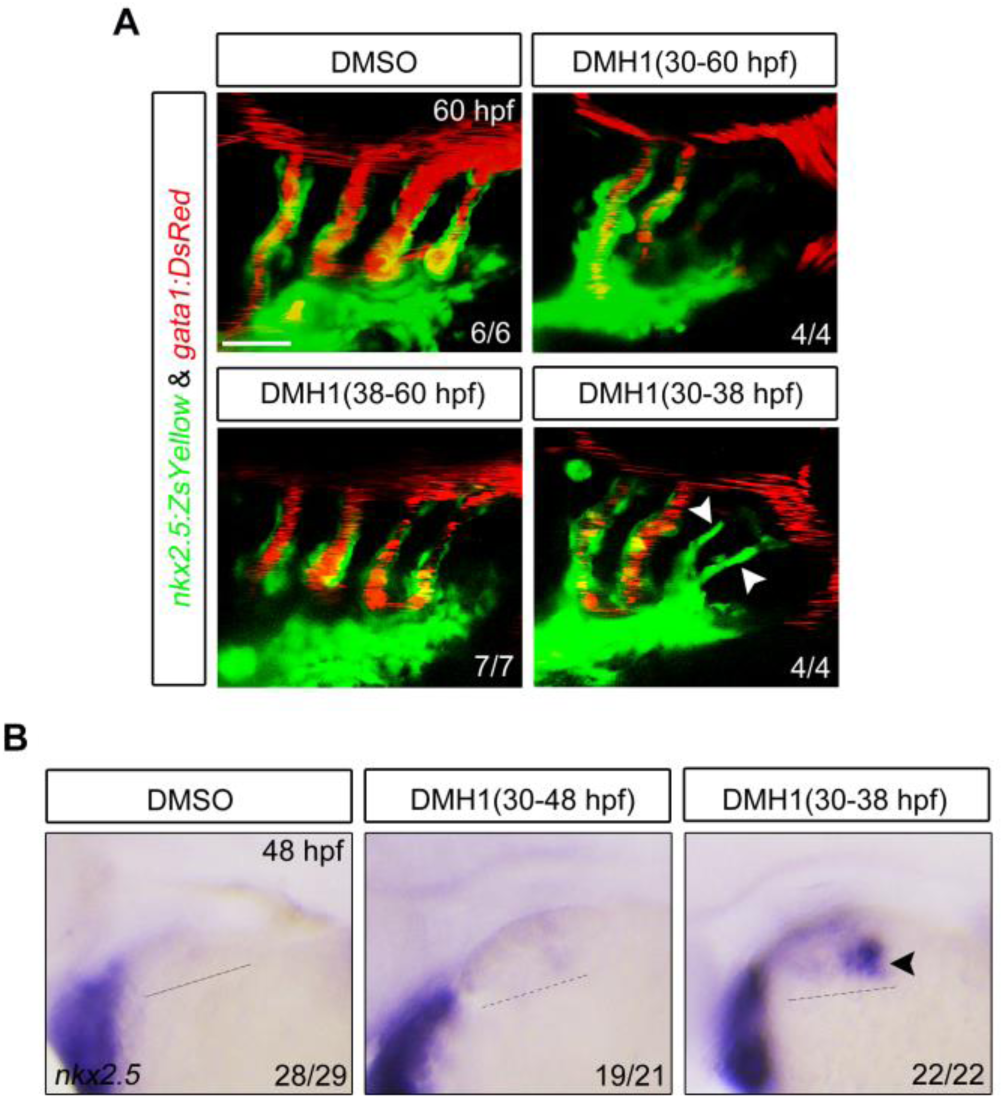
Blockage of BMP signaling after PAA progenitor specification induces no defects of PAA morphogenesis. (A) Live confocal images of *Tg(nkx2.5:ZsYellow;gata1:DsRed)* embryos at 60 hpf with DMSO or 10 μM DMH1 treatment for different durations. White arrowheads indicate the new sprouting clusters after removal of DMH1 at 38 hpf. Scale bar, 50 μm. (B) Expression analysis of *nkx2.5* by in situ hybridization in embryos subjected to different treatments. The dotted lines represent the area of expression of *nkx2.5*. Black arrowhead indicates the newly emerged expression of *nkx2.5* after removal of DMH1.

### BMP2a and BMP5 have redundant functions in PAA progenitor specification

To explore which BMP ligands are specifically required for PAA progenitor specialization, knockdown experiments were performed using antisense MOs that had previously been used to interfere the expression of *bmp2a*, *bmp2b*, *bmp4*, and *bmp5* (Chocron et al., 2007; Li et al., 2019; Naye et al., 2012; Shih et al., 2017). As expected, injection of these MOs into wild-type embryos caused clear defects in the induction of the hepatic bud, the formation of pharyngeal pouches, the left-right asymmetry of the embryonic heart, and the development of neural crest cells, respectively (S7A-S7E Fig) (Chocron et al., 2007; Li et al., 2019; Naye et al., 2012; Shih et al., 2017), indicating a satisfactory level of efficiency and specificity of these MOs. We observed that the expression of PAA angioblast marker *etv2* was not obviously changed in embryos injected with *bmp4* MO (Fig 6A and 6B). It was surprising that the expression of *etv2* was almost abolished in *bmp2b* morphants (Fig 6A and 6B). We have previously reported that *bmp2b* is essential for pharyngeal pouch progenitor specification (Li et al., 2019). In fact, compared to control embryos, *bmp2b* morphants showed no pharyngeal pouches at 36 hpf, as indicated by the expression of pouch epithelium marker *nkx2.3* (S7B Fig). Therefore, although we cannot rule out that *bmp2b* plays a direct role in PAA development, the loss of PAA angioblast in *bmp2b* morphants is due mainly to the deficiency of pharyngeal pouches. Importantly, knockdown of *bmp2a* or *bmp5* resulted in a steady reduction in *etv2^+^* cluster numbers, and this reduction became more pronounced when these two genes were knocked down at the same time (Fig 6A and 6B). Furthermore, the phosphorylation of Smad1/5/8 and the expression of *nkx2.5* were evidently decreased in the pharyngeal region (Fig 6C and 6D). Together, these data suggest that *bmp2a* together with *bmp5* plays an important role in PAA progenitor specification.

**Fig 6.**
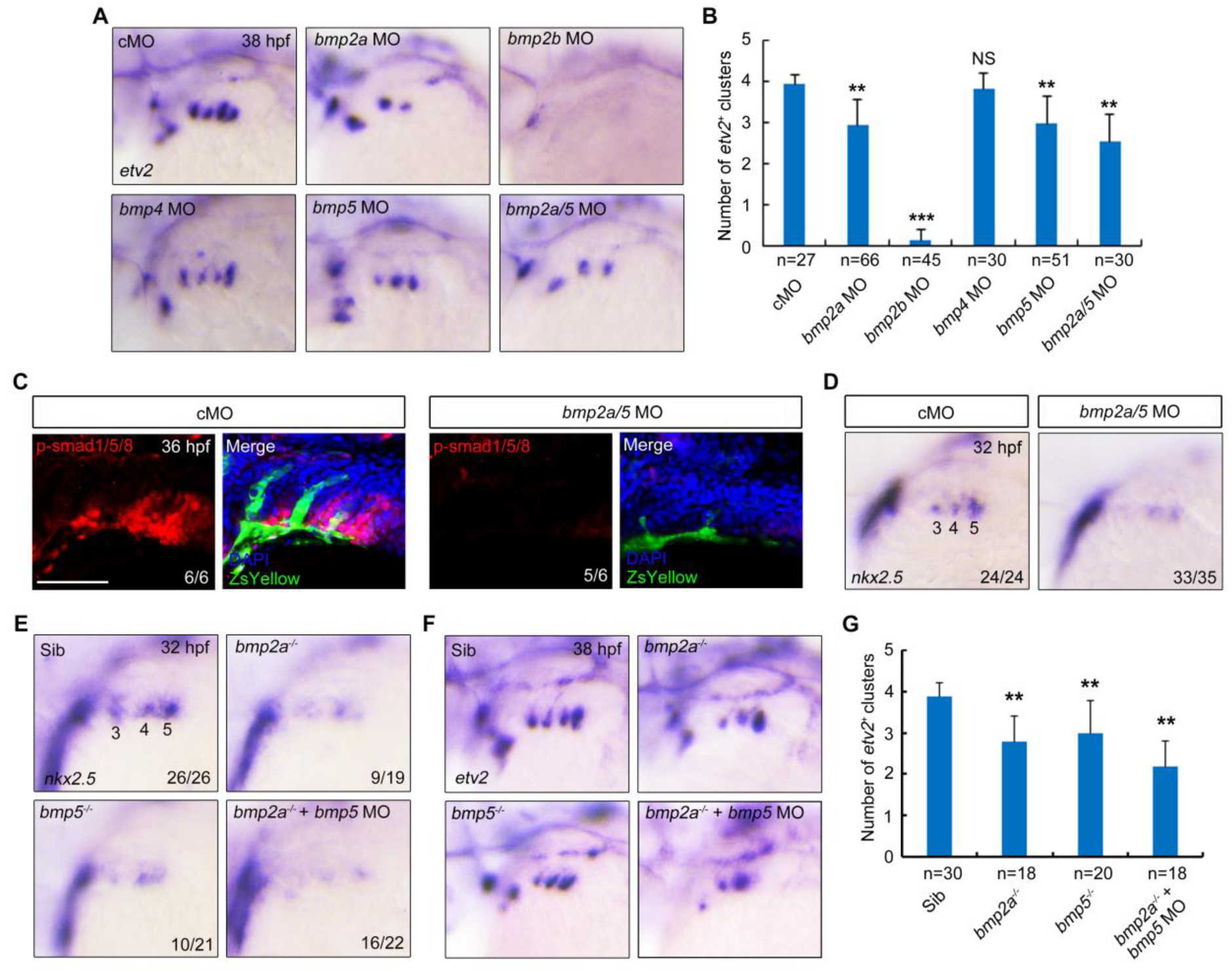
BMP2a together with BMP5 functions in PAA progenitor specification. (A-B) The expression of *etv2* in embryos injected with indicated MO was analyzed by *in situ* hybridization (A). Injection doses: *bmp2a* MO, 2 ng; *bmp2b* MO, 0.3 ng; *bmp4* MO, 2ng; *bmp5* MO, 4ng. The average numbers of *etv2^+^* PAA angioblast clusters were quantified from three independent experiments, and the group values were expressed as mean ± SD (B). Student’s *t*-test, \*\**p* < 0.01. NS, no significant difference. (C-D) Knockdown of *bmp2a* and *bmp5* restrained BMP signal activation in the pharyngeal region and disrupted the specification of PAA progenitors. *Tg(nkx2.5:ZsYellow)* or wild-type embryos were injected with *bmp2a* and *bmp5* MOs at the one-cell stage. The resulting embryos were harvested at indicated stages for immunostaining (C) or *in situ* hybridization (D). Scale bar, 50 μm. (E-G) Expression analysis of *nkx2.5* (E) and *etv2* (F) in *bmp2a^-/-^* or *bmp5^-/-^* single mutants and *bmp2a^-/-^* mutant embryos injected with 4 ng *bmp5* MO by *in situ* hybridization. The average numbers of *etv2^+^* angioblast clusters were quantified from three independent experiments and the group values were expressed as mean ± SD (G). Student’s *t*-test, \*\**p* < 0.01. Note that depletion of *bmp2a* or/and *bmp5* suppressed the specification and angioblast differentiation of PAA progenitors.

To examine the direct function of pouch-expressed BMP ligands, we performed tissue-specific knockdown experiments using a KalTA4-UAS system to drive the expression of miR30-based short hairpin RNAs (shRNAs), which is widely used for efficient gene silencing in eukaryotic organisms (S8A Fig) (Li et al., 2018; Stegmeier et al., 2005; Zeng et al., 2005). We first generated two *UAS:EGFP-shRNA* plasmids expressing shRNAs targeting different sequences of *bmp2a* (named as *shRNA-bmp2a-1* and *shRNA-bmp2a-2*). *KalTA4-p2a-mCherry* mRNA, together with *shRNA-bmp2a-1* or *shRNA-bmp2a-2,* were co-injected into one-cell stage embryos, and the expression of *bmp2a* was examined by WISH at 11 hpf. As shown in S6B Figure, KalTA4-mediated expression of *shRNA-bmp2a-2* clearly knocked down endogenous *bmp2a* expression. Similarly, we found that *shRNA-bmp2b-1*, *shRNA-bmp4-2* and *shRNA-bmp5-1*, could effectively silence genes when employed independently (S8C-S8E Fig).

Next, these selected *UAS:EGFP-shRNA* plasmids and Tol2 transposase mRNA were injected into *Tg(nkx2.3:KalTA4-p2a-mCherry)* embryos. A subset of the resulting embryos showed bright green fluorescence in the pharyngeal pouches at 36 hpf as previously reported (S8F Fig) (Li et al., 2018). Such embryos were collected to examine the developmental consequences of *bmp* gene deficiency. Although the depletion of *bmp2b* or *bmp4* expression in pouches demonstrated no effect on angioblast formation, silencing of *bmp2a* or *bmp5* eliminated the expression of *etv2* in the pharyngeal region (S8G and S8H Fig). In addition, attenuation of the function of both *bmp2a* and *bmp5* resulted in a more evident decline in the number of PAA angioblast clusters (S8G and S8H Fig). Besides, simultaneous knockdown of *bmp2a* and *bmp5* led to severe decrease in the phosphorylation of Smad1/5/8 in the pharyngeal region and reduced expression of *nkx2.5* in the PAA clusters (S8I and S8J Fig). These findings support the idea that pouch-derived BMP2a and BMP5, not BMP2b and BMP4, are responsible for PAA progenitor specification.

To further substantiate the function of *bmp2a* and *bmp5* in PAA development, we generated one genetic mutant line for each gene using CRISPR/CAS9 technology. The mutant allele of *bmp2a* or *bmp5* carries a DNA deletion near the gRNA targeting sequence in the first exon, resulting in a premature stop codon and presumably a truncated protein lacking the prodomain and C-terminal mature peptide (S9A and S9B Fig). *In situ* hybridization results revealed that about 50% of *bmp2a^-/-^* mutants, which were confirmed by genotyping, showed a decrease in *hhex* expression in the hepatic bud at 30 hpf (S9C Fig). A similar percentage of *bmp5^-/-^* mutant embryos with defective development of neural crest cells was observed at 24 hpf (S9D Fig). Previous studies have suggested that a genetic mutation may induce an increase of related gene expression to assume the function of the mutated gene (El-Brolosy et al., 2019; Rossi et al., 2015). Thus, the incomplete penetrance in *bmp2a* and *bmp5* mutants may be due to a certain degree of compensatory adaptations. Consistent with this idea, we observed that only approximately half of *bmp2a^-/-^* and *bmp5^-/-^* embryos exhibited impaired formation of PAA progenitors at 32 hpf (Fig 6E). By contrast, injection of *bmp5* MO into to *bmp2a^-/-^* embryos induced more obvious defects in PAA progenitor specification in a much higher proportion of animals (Fig 6E). Finally, at 38 hpf, a significant reduction in *etv2*^+^ angioblast clusters was observed in *bmp2a^-/-^* mutants injected with *bmp5* MO when compared to control animals and *bmp2a^-/-^* or *bmp5^-/-^* embryos (Fig 6F and 6G).

Collectively, these data suggest that, among the BMPs emerged from the pouch endoderm, BMP2a and BMP5 are crucial for BMP pathway activation in the pharyngeal mesoderm, thereby promoting PAA progenitor specification.

### *npas4l* is essential for endothelial lineage progression from the pharyngeal mesoderm to PAA progenitors

The progenitors for PAA, HM and cardiac OFT are situated within the pharyngeal mesoderm and are both marked by *nkx2.5* expression (Guner-Ataman et al., 2018; Paffett-Lugassy et al., 2013). Future identification of specific biomolecular markers for PAA progenitors can provide new avenues to investigate the cellular and molecular events in PAA progenitor specification. A recent study identified *npas4l*, which encodes a PAS-domain-containing bHLH transcription factor, as the gene defective in the well-known *cloche* mutant that lacks most endothelial as well as hematopoietic cells (Reischauer et al., 2016). Therefore, we speculate that *npas4l* may be expressed in PAA progenitors and critical for PAA development. To verify this hypothesis, the expression of *npas4l* in the pharyngeal region was analyzed by *in situ* hybridization. We found that *npas4l* was not expressed in the pharynx at 20 hpf (Fig 7A). But then *npas4l* transcripts was detected in the presumptive PAA progenitor cluster 3 at 24 hpf, approximately 2 hours later than the initial expression of *nkx2.5* in the same PAA cluster (Fig 7A). Over the next 14 hours, *npas4l* transcripts were gradually appeared in a craniocaudal sequence in the PAA clusters (Fig 7 B). Moreover, the expression of *npas4l* in the PAA clusters was further confirmed by the colocation of *npas4l* and *nkx2.5* transcripts (Fig 7C).

**Fig 7.**
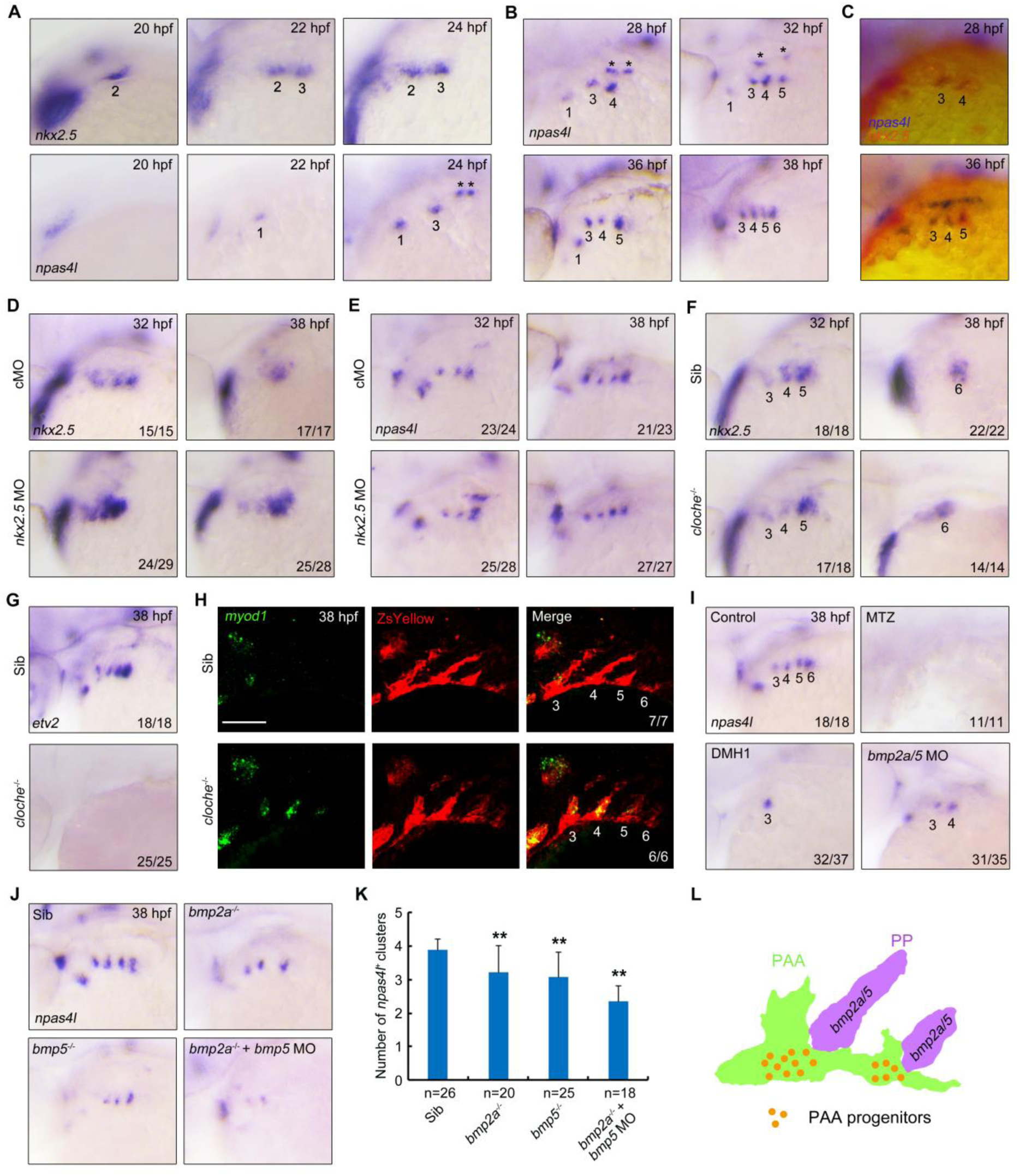
*npas4l* plays a pivotal role in the specification of PAA progenitors from pharyngeal mesoderm. (A-B) Analysis the expression patterns of *nkx2.5* and *npas4l* in the pharynx at the indicated stages by *in situ* hybridization. Lateral view, head towards the left. The asterisk indicates the expression of *npas4l* in the LDA. (C) Double *in situ* hybridization of *npas4l* (red) and *nkx2.5* (blue) expression at 28 and 38 hpf. (D-E) Expression analysis of *nkx2.5* (D) and *npas4l* (E) in embryos injected with 3 ng *nkx2.5* MO. Note that the expression of *nkx2.5* but not *npas4l* was clearly increased in *nkx2.5* morphants. (F-G) The expression of *nkx2.5* (F) and *etv2* (G) were analyzed in *cloche^-/-^* mutants and their wild-type and heterozygous siblings by *in situ* hybridization. (H) The expression of *myod1* in the pharynx was analyzed in *cloche^-/-^* mutants and their siblings in *Tg(nkx2.5:ZsYellow)* background. The embryos were firstly subjected to fluorescent *in situ* hybridization with *myod1* probe (green), and then stained with ZsYellow antibody (red). (I) *Tg(nkx2.3:KalTA4-p2a-mCherry;UAS:NTR-mCherry)* embryos were treated with 10 mM MTZ from bud stage and wild-type embryos were treated with 10 μM DMH1 from 24 hpf or injected with 2 ng *bmp2a* MO together with 4 ng *bmp5* MO at the one-cell stage. The resulting embryos were harvested for *in situ* hybridization with *npas4l* probe at 38 hpf. (J-K) Expression analysis of *npas4l* in *bmp2a^-/-^* or *bmp5^-/-^* single mutants and *bmp2a^-/-^* mutant embryos injected with 2ng *bmp5* MO (J). The average numbers of *npas4l^+^* clusters were quantified from three independent experiments and the group values were expressed as mean ± SD (K). Student’s *t*-test, \*\**p* < 0.01. (L) Working model depicts that the specification of PAA progenitors from pharyngeal mesoderm is dependent of the activation of BMP signaling by *bmp2a* and *bmp5* expressed in pouch endoderm.

Previous study has shown that the expression of *nkx2.5* is reduced following the differentiation of PAA progenitors into angioblasts (Paffett-Lugassy et al., 2013). Intriguingly, *npas4l* transcripts persisted during PAA progenitor differentiation (Fig 7B). These observations raised a possibility that *npas4l* is not only expressed in the progenitor but also in the angioblasts of PAAs. Previous study has suggested that the disruption of angioblast differentiation would result in an accumulation of PAA progenitors (Abrial et al., 2017; Paffett-Lugassy et al., 2013). Because *nkx2.5* is not required for PAA progenitor specification but essential for angioblast differentiation (Paffett-Lugassy et al., 2013), we then examined the expression of *npas4l* in *nkx2.5* morphants. If *npas4l* is specifically expressed in PAA progenitors, we would expect a clear increase of *npas4l* expression in *nkx2.5* morphants. Indeed, compared to control animals, embryos injected with *nkx2.5* MO showed much higher levels of *nkx2.5* expression (Fig 7D). By contrast, the expression levels of *npas4l* was not obviously changed in the pharynx upon *nkx2.5* MO injection (Fig 7E). These results may imply that although the inhibition of *nkx2.5* function led to excess PAA progenitors at the expense of angioblasts, the total number of cells with endothelial potential was unchanged. Thus, *npas4l* is expressed in both PAA progenitors and angioblasts.

We next examined whether *npas4l* plays a role in PAA development. *In situ hybridization* analyses revealed that, in comparison to wild-type and heterozygous siblings, *cloche* homozygous (*cloche^-/-^)* mutants exhibited normal *nkx2.5* expression in pharyngeal clusters 3-5 at 32 hpf (Fig 7F). To our surprise, the expression of *etv2*, the PAA angioblast marker, was completely missing in the *cloche^-/-^* mutants at 38 hpf (Fig 7G), suggesting a successful differentiation of *nkx2.5^+^* progenitors. This phenomenon thus raised an interesting question about the cell fate of the *nkx2.5^+^* progenitors in *cloche^-/-^* mutants. It has been suggested that the *nkx2.5^+^* progenitors in lateral plate mesoderm can differentiate into various pharyngeal tissues including PAA, HM and cardiac OFT (Guner-Ataman et al., 2018; Paffett-Lugassy et al., 2013). Therefore, we investigated whether the *nkx2.5^+^* progenitors within presumptive PAA clusters in *cloche^-/-^* mutants altered their cell fate to give rise to cardiac OFT and/or to become muscle cells. We found no distinct difference in the expression of *ltbp3*, which labels the outflow pole of the heart tube (Zhou et al., 2011), between *cloche^-/-^* mutants and their siblings (S10 Fig). On the contrary, the transcripts of *myod1*, the head muscle precursor marker (Lin et al., 2006), were unexpectedly expressed in the presumptive PAA clusters of *cloche^-/-^* mutants (Fig 7H), indicating a muscle cell fate transformation of the *nkx2.5^+^* progenitors. Therefore, before *npas4l* expression, the pharyngeal mesoderm might maintain its multiple differentiation potential into at least the endothelial and muscular lineages. Together, these data suggest that *npas4l* plays a pivotal role in the specification of PAA progenitors from pharyngeal mesoderm.

To learn whether pharyngeal pouches are required for *npas4l* expression, *Tg(nkx2.3:KalTA4-p2a-mCherry;UAS:NTR-mCherry)* embryos were exposed to MTZ from the bud stage. Ablation of pouch endoderm completely abolished *npas4l* expression in the pharynx at 38 hpf (Fig 7I). Moreover, both DMH1 treatment and injection with MOs targeting *bmp2a* and *bmp5* induced a dramatic reduction in *npas4l* transcripts (Fig 7I). We also found a steady decrease of the number of *npas4l^+^* PAA clusters in *bmp2a^-/-^* or *bmp5^-/-^* single mutants and *bmp2a^-/-^* embryos injected with *bmp5* MO (Fig 7J and 7K). Taken together, these findings support the idea that the pharyngeal pouches provide a niche microenvironment for the commitment of multipotent pharyngeal mesoderm toward PAA progenitors through expressing BMP2a and BMP5 (Fig. 7L).

## Discussion

Improper embryonic development of the PAAs may cause life-threating congenital cardiovascular defects (Abrial et al., 2017; Srivastava, 2001). Malformations of the aortic arch system were often accompanied by anomalies of endodermal pouches, which would lead to compromised pharyngeal segmentation (Jerome and Papaioannou, 2001; Lindsay et al., 2001; Piotrowski et al., 2003). The effects of pouches on aortic arch development have traditionally been considered secondary to pharyngeal patterning defects (Kopinke et al., 2006; Matt et al., 2003; Wendling et al., 2000). In this study, our results support a model in which the pharyngeal pouches provide a niche microenvironment for PAA progenitor specification via the expression of BMP proteins. Our findings suggest that the segmentation of pharyngeal pouches coincides spatiotemporally with the emergence of PAA progenitor clusters. Furthermore, depletion of pouch endoderm in zebrafish embryos by a MTZ-NTR system resulted in a remarkable reduction of BMP signal activity in the pharyngeal mesoderm and the complete absence of PAA structures attributed to impaired progenitor specification.

Most importantly, the PAA progenitor specification is directly regulated by pouches and their derived signal molecules, as the ablation of pharyngeal neural crest cells shows no effect on the emergence of PAA angioblasts, which are differentiated from vascular progenitors. These data, combined with our recent findings that pharyngeal pouches regulate PAA angioblast proliferation by expressing PDGF ligands (Mao et al., 2019), suggest multiple distinct roles of pouch endoderm during PAA development.

It has been reported that a common progenitor population for PAAs, HM and cardiac OFT is specified in zebrafish ALPM during mid-somitogenesis (Guner-Ataman et al., 2018). At later stages, these progenitors located within pharyngeal arches 3-6 contribute to the endothelium of their respective PAAs (Paffett-Lugassy et al., 2013). Interestingly, our data indicate that, at later stages, this pharyngeal mesoderm lineage within PAs 3-6 is comprised of two subpopulations: one of which is *nkx2.5^+^* which are cells restricted at different domains, and the other is *nkx2.5^-^* which are cells located between the *nkx2.5^+^* clusters. Consistent with previous findings, our cell-lineage tracing analysis reveals that the *nkx2.5^+^* clusters sprout out and contribute to corresponding PAAs (Paffett-Lugassy et al., 2013). Previous reports have also suggested that, in zebrafish, the ventral parts of PAAs merge to form the bilateral ventral aortae (Anderson et al., 2008; Paffett-Lugassy et al., 2013). Unexpectedly, we found that the *nkx2.5^-^* subpopulation does not migrate dorsally and ultimately gives rise to ventral aortae. Therefore, PAAs are sequentially generated from *nkx2.5^+^* progenitors within the developing ventral aortae (the pharyngeal mesoderm). Interestingly, resin filling of mouse embryonic vasculature has shown that the PAA endothelium arises by branching off from the aortic sac, the mammalian homolog of the ventral aorta of gill-bearing vertebrates (Annalisa Berta, 2006; Hiruma et al., 2002). Based on these results, it is likely that the process of PAA morphogenesis is evolutionarily conserved across vertebrate classes.

Several lines of evidence support the idea that the pharyngeal mesoderm lineage is further specified into PAA progenitors in a niche microenvironment provided by pouches. Firstly, the pharyngeal mesoderm within PAs 3-6 contains *nkx2.5^+^* progenitors that give rise to PAAs, and *nkx2.5^-^* progenitors that generate ventral aorta. Secondly, the PAA progenitor clusters emerge in a craniocaudal sequence following pharyngeal pouch segmentation. Thirdly, depletion of pouch endoderm eliminates the PAA progenitors without disrupting the segregation of pharyngeal mesoderm lineage from cardiac precursors. Fourthly, BMP signal inhibition by pharmacological treatments and tissue-specific knockdown or genetic depletion of *bmp2a*/5 results in remarkably reduced PAA progenitors. Finally and yet most importantly, *cloche^-/-^* mutants exhibit ectopic head muscle progenitors in the presumptive PAA clusters, indicating that the pharyngeal mesoderm might have multiple differentiation potential before the expression of the *cloche* gene *npas4l*, which is gradually appeared in PAA progenitors but not in HM and cardiac OFT precursors. This idea is supported by a previous observation that in *cloche^-/-^* mutants, the rostral mesoderm undergoes a fate transformation and generates ectopic cardiomyocytes (Schoenebeck et al., 2007).

Our study demonstrates that *npas4l* is essential for PAA progenitor specification and its expression is tightly controlled by BMP signaling. Moreover, mutation of the *cloche* locus result in a cell fate transformation: rather than producing progenitors of PAAs, the pharyngeal mesoderm produces ectopic head muscle progenitors. Interestingly, when BMP signal inhibition is relieved, PAA progenitors are capable of reappearing in the pharyngeal mesoderm and develop into growing sprouts. Previous studies have reveal that *npas4l* functions upstream of *etv2*, and overexpression of *etv2* induces vascular gene expression and converts skeletal muscle cells into functional endothelial cells (Reischauer et al., 2016; Veldman et al., 2013). Therefore, when BMP signal is reactivated after the washout of BMP inhibitors, the expression of *npas4l* and its downstream gene *etv2* might be reinduced in the muscle progenitors located in the presumptive PAA clusters and then transdifferentiate them into endothelial cells. Additional studies will be required to learn whether BMP-Npas4l-Etv2 pathway is necessary and sufficient to switch the fate of muscle cells into the vascular lineage in the pharyngeal region.

During vertebrate embryonic development, pharyngeal pouches play a central role in organization of the head through expressing signaling molecules like FGFs and BMPs (Crump et al., 2004; David et al., 2002; Graham, 2008; Holzschuh et al., 2005; Ning et al., 2013). Our data further indicate that pharyngeal pouches induce PAA progenitors by expressing BMP ligands. Stem cells or progenitor populations are established in niches where niche factors function to maintain their quiescent state or to induce their proliferation and differentiation for fetal development (Birbrair and Frenette, 2016; Jhala and Vasita, 2015; Scadden, 2006). Since endoderm pouches are in close contact with the pharyngeal mesoderm at discrete locations, establishing a physicochemical environment for cell fate determination through the activation of BMP signaling, it is reasonable to hypothesize that pharyngeal pouches provide a niche microenvironment for PAA progenitor specification.

Genetic inactivation of various components of the canonical BMP signaling cascade results in abnormal vascular remodeling of the PAAs (Liu et al., 2004; Nie et al., 2008; Ohnemus et al., 2002; Papangeli and Scambler, 2013). Using the KalTA4-UAS system to drive pouch-specific expression of miR30-based short hairpin RNAs, we find that both *bmp2a* and *bmp5* are responsible for progenitor specification. We speculate that the redundant functions of these *bmp* genes may be responsible for impeding the discovery of their potential roles in PAA progenitor commitment in previous studies. In fact, pouch endoderm expresses several *bmp* genes including *bmp2a*, *bmp2b*, *bmp4* and *bmp7*. Genetic interference approaches have identified that *bmp2b* and *bmp5* function redundantly in epibranchial neurogenesis (Holzschuh et al., 2005). Since different BMP ligands activate signaling pathways through different receptors, it is possible that the distinct expression profiles of BMP receptors in the pharyngeal mesoderm plays a role in shaping the requirement for BMP5 together with BMP2a, but not for other BMPs, in PAA progenitor specification.

### Materials and methods Ethics statement

Our zebrafish experiments were all approved and carried out in accordance with the Animal Care Committee at the Institute of Zoology, Chinese Academy of Sciences (Permission Number: IOZ-13048).

### Zebrafish lines

Our zebrafish experiments were performed by using the following mutant and transgenic lines: *Tg*(*nkx2.5:ZsYellow*) (Paffett-Lugassy et al., 2013), *Tg*(*nkx2.5:kaede*) (Paffett-Lugassy et al., 2013), *Tg*(*nkx2.3:mCherry*) (Li et al., 2018), *Tg*(*nkx2.3:KalTA4-p2a-mCherry*) (Li et al., 2018), *Tg*(*flk:EGFP*), *Tg*(*gata1:DsRed*), *Tg*(*sox17:GFP*), *Tg*(*sclβ:d2eGFP*) (Zhen et al., 2013), *Tg*(*BRE:EGFP*) (Laux et al., 2011), *Tg(hsp70l:dnBmpr1a-GFP) (Pyati et al., 2005)*, *Tg(hsp70l:caBmpr1b-GFP) (Row and Kimelman, 2009)*, *Tg*(*sox10:KalTA4-p2a-mCherry*), *Tg(UAS:NTR-mCherry)* and *cloche*^m378^ (Stainier et al., 1995). *Tg*(*sox10:KalTA4-p2a-mCherry*) transgenic line was generated by our lab with the *sox10* upstream regulatory sequence as previously described (Carney et al., 2006). *Tg(UAS:NTR-mCherry)* transgenic line was obtained from China Zebrafish Resource Center. Unless otherwise specified, live embryos were kept at 28.5 °C in Holtfreter’s solution, and staged based on morphology as previously described (Kimmel et al., 1995).

### RNA synthesis and WISH

Capped RNAs for casmad1 and casmad5 were synthesized *in vitro* from corresponding linearized plasmids using the mMessage mMachine kit (Ambion). Digoxigenin-UTP-labelled antisense RNA probes for *scl*, *etv2*, *nkx2.5*, *ZsYellow*, *tie1*, *sox17*, *bmp2a*, *bmp2b*, *bmp4, bmp5* and *npas4l* were transcribed using MEGAscript Kit (Ambion) according to the manufacturer’s instructions. WISH with these RNA probes were performed using the NBT-BCIP substrate.

### Morpholinos and microinjections

Morpholino oligonucleotides (MOs) were purchased from Gene Tools (Philomath, OR, USA). The standard control MO (5’-CCTCTTACCTCAGTTACAATTTATA-3’) (Dickmeis et al., 2001), *sox32* MO (5’-CAGGGAGCATCCGGTCGAGATACAT-3’) (Dickmeis et al., 2001), *bmp2a* MO (5’-AGTAAACACTTGCTTACCATCATGG-3’) (Naye et al., 2012), *bmp2b* MO (5’-CGCGGACCACGGCGACCATGATC-3’) (Li et al., 2019), *bmp4* MO (5’-GTCTCGACAGAAAATAAAGCATGGG-3’) (Chocron et al., 2007), *bmp5* MO (5’- TTGACCAGGATGATGATGCTTTCAG-3’) (Shih et al., 2017) and *nkx2.5* MO (5’-TGTCAAGGCTCACCTTTTTTCTCTT-3’) (Paffett-Lugassy et al., 2013) were used as previously described. All the MOs were injected at one-cell stage in zebrafish embryos.

### MiR30-based shRNAs

The miR30-based shRNAs were designed according to previously published methods (Dow et al., 2012). The target sequences were shown in S1 Table. Plasmids expressing shRNAs were microinjected into fertilized eggs at the one-cell stage at indicated concentrations. The injected embryos were cultured at 28.5 °C till further operation.

### Generation of *bmp2a* and *bmp5* mutants

We generated *bmp2a* and *bmp5* mutants using the CRISPR/CAS9 technology. We designed the gRNAs of *bmp2a* and *bmp5* using the CRISPR Design website http://crispor.tefor.net/. The Cas9 mRNA and gRNAs were prepared as described before (Wei et al., 2017), and co-injected into wild-type embryos at the one-cell stage. Embryos were collected to make genomic DNA for genotyping at 24 hpf. For screening of the F1 fish with mutant alleles, genomic DNA was isolated from the tail of individual fish. The forward primer 5’-AAAGACTCGCAATGGCTCG-3’ and reverse primer 5’-TCCCTGTCAGGCATGAAG-3’ were used to amplify *bmp2a* gRNA targeted sequence. And the forward primer 5’-GACTTCTGTGGAGCTGTTTAG-3’ and reverse primer 5’-TGCGTGACCTCTTTACACCAT-3’ were used to amplify *bmp5* gRNA targeted sequence. The amplified fragments were identified with Sanger DNA sequencing for genotyping. F2 embryos were generated by incrossing F1 mutant fishes and genotyped by digesting PCR products with BtsI (NEB, R0667S) and SmaI (NEB, R0141V), respectively.

### Live embryo imaging and kaede photoconversion

Live fluorescent embryos were mounted in 1% low-melting agarose in glass bottom dish (Solarbio; D35-10-1-N) at indicated stages. *Tg*(*nkx2.5: ZsYellow*) embryos were imaged and analyzed for the formation of PAAs using a Nikon A1R+ confocal microscope (20× objective). For cell lineage trancing, photoconversion in *Tg(nkx2.5:kaede)* embryos was achieved by DAPI filter, and the converted embryos were immediately imaged, removed from the agarose, and raised in dark conditions until subsequent evaluation. All confocal stack images were processed using the Nikon NIS-Elements AR 4.13.00 software.

### Immunofluorescence staining and fluorescent *in situ* hybridization

Immunofluorescence staining was performed as previously described (Ning et al., 2013). Embryos were stained with the following affinity-purified antibodies: anti-GFP (1:1000; A111201, Invitrogen), anti-ZsYellow (1:200; 632475, Clontech), anti-ZsYellow (1:400; TA180004, Origene), anti-p-Smad1/5/8 (1:200; 9511, Cell Signaling Technology). Fluorescence *in situ* hybridization was performed as previously described (Schoenebeck et al., 2007). Anti-DIG HRP-conjugated Fab fragments (1:400; Roche) were used to detect the digoxigenin (DIG)-labeled probes. Then, embryos were incubated with fluorescein (FLU) tyramide (1:100; PerkinElmer) for 3 hours at 28.5 °C. Next, the embryos were subjected to immunofluorescence after removal of HRP activity.

### Pharmacological treatment and heat shock

For tissue-specific ablation, *Tg(nkx2.3:KalTA4-p2a-mCherry;UAS:NTR-mCherry)* or *Tg(sox10:KalTA4-p2a-mCherry;UAS:NTR-mCherry)* embryos were raised in Holtfreter’s solution containing 10 mM MTZ (M1547, Sigma) from bud stage, and then harvested for live imaging or *in situ* hybridizations at the indicated stages. To block BMP signaling, embryos were treated with DMH1 (10 μM; D8946, Sigma) or Dorsomorphin (10 μM; P5499, Sigma) under dark conditions. *Tg(hsp70l:dnBmpr1a-GFP)* and *Tg(hsp70l:caBmpr1b-GFP)* embryos were subjected to heat shock at 40 °C for 20 min at 24 hpf, and then incubated at 28.5 ℃ until harvest.

## Statistical analysis

Statistical analysis was performed with GraphPad Prism software version 5.00 for Macintosh (GraphPad). The numbers of PAA angioblast clusters were counted based on PAA3-PAA6 per embryo. All results were expressed as mean ± SD. Differences between control and treated groups were analyzed with unpaired two-tailed Student’s *t*-test. Results were considered statistically significant at p <0.05.

## Acknowledgements

We are grateful to Dr. Jingwei Xiong (Peking University, China) for *Tg*(*nkx2.5:ZsYellow*) fish line and Dr. Caroline E. Burns (Massachusetts General Hospital, USA) for *Tg(nkx2.5:kaede)* fish line.

## Supporting information

### Supplemental Figures

**S1 Fig.**
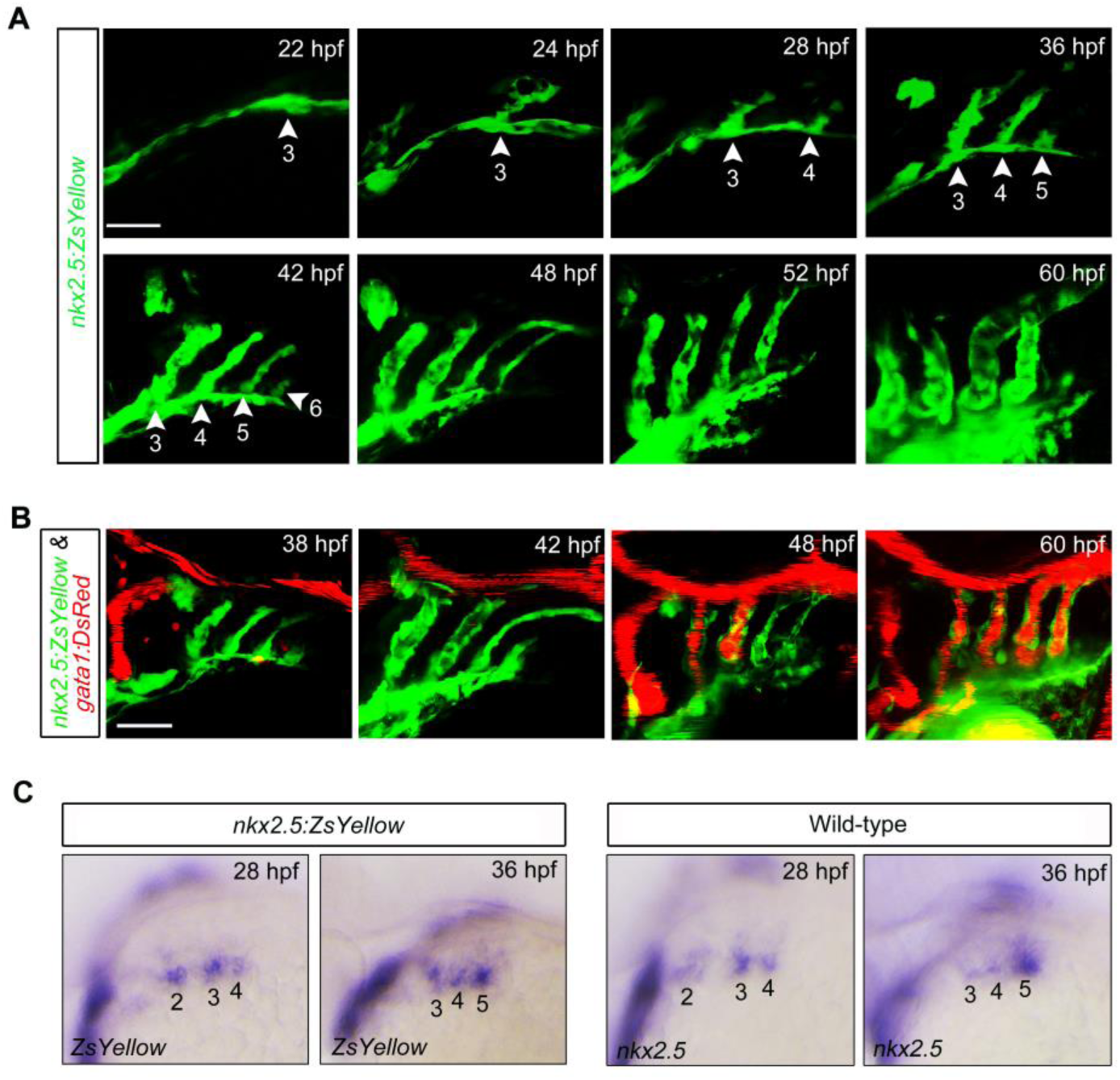
The dynamic establishment of the functional aortic arches. (A) Time-lapse recording of ZsYellow fluorescence in the pharyngeal region of *Tg(nkx2.5:ZsYellow)* embryos from 22 to 60 hpf shows the sequential emergence of PAA clusters and the step-by-step establishment of PAAs. White arrowheads highlight the forming PAA clusters or the PAA sprouts, which are numbered with Arabic numerals. All the embryos are shown in lateral views with anterior on the left. Scale bar, 50 μm. (B) Live confocal images of *Tg(nkx2.5:ZsYellow;gata1:DsRed)* embryos with green endothelial cells and red erythrocytes at indicated developmental stages. All the embryos were shown in lateral views with anterior on the left. Scale bar, 50 μm. (C) *In situ* hybridization analysis of the expression patterns of *ZsYellow* and *nkx2.5* in the pharynx of *Tg(nkx2.5:ZsYellow)* or wild-type embryos at 28 and 36 hpf. Numbers indicate PAA clusters. Lateral views with anterior on the left.

**S2 Fig.**
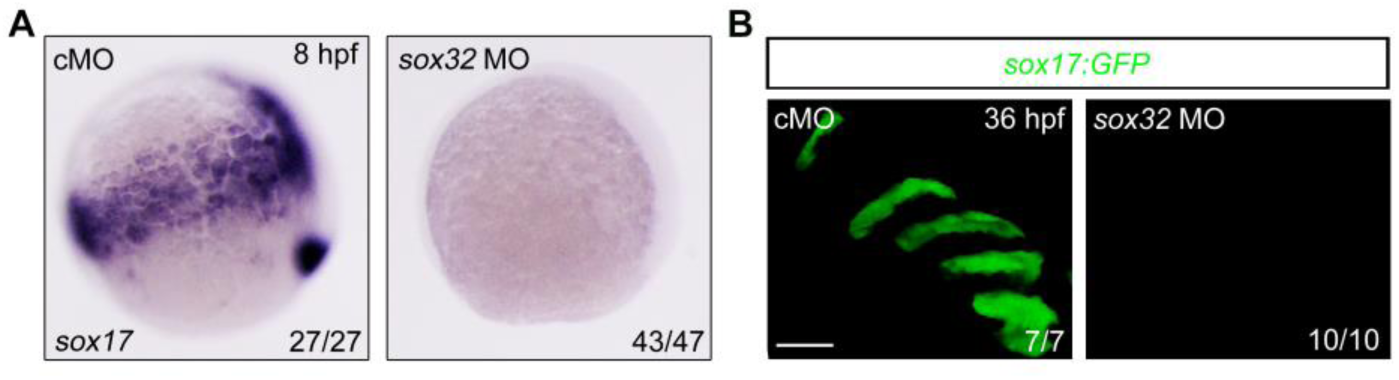
Injection of *sox32* MO results in absence of endodermal cells and pharyngeal pouches. (A) Expression analysis of *sox17* by *in situ* hybridization at 8 hpf in embryos injected with 8 ng cMO or *sox32* MO. (B) Live images of pharyngeal pouches in *Tg(sox17:GFP)* embryos at 36 hpf, which were injected with 8 ng cMO or *sox32* MO. Scale bar, 50 μm.

**S3 Fig.**
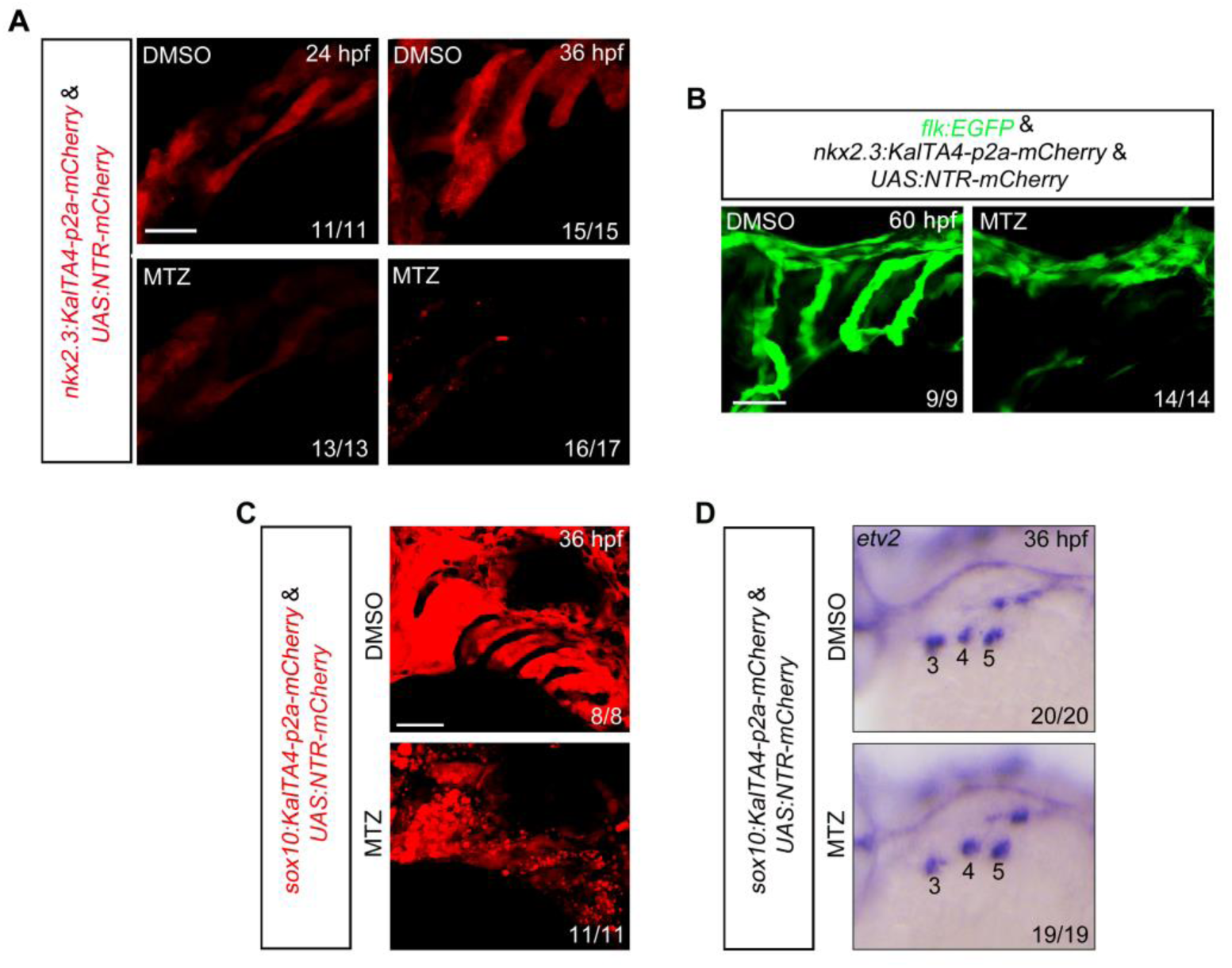
Tissue ablation by MTZ-NTR system. (A) *Tg(nkx2.3:KalTA4-p2a-mCherry;UAS:NTR-mCherry)* embryos were treated with 10 mM MTZ from bud stage. These embryos were harvested at 24 or 36 hpf for *in vivo* confocal imaging. The mCherry fluorescence was greatly reduced in MTZ-treated embryos. Scale bar, 50 μm. (B) Embryos were exposed to 10 mM MTZ from bud stage to 38 hpf, and then harvested at 60 hpf for *in vivo* confocal imaging. Scale bar, 50 μm. (C-D) *Tg(sox10:KalTA4-p2a-mCherry;UAS:NTR-mCherry)* embryos were treated with 10 mM MTZ from bud stage to 36 hpf, and then harvested for *in vivo* confocal imaging (C) and *in situ* hybridizations (D). Note that the MTZ treatment induced evident cell death in pharyngeal neural crest cells, but showed no effect on PAA angioblast development. Scale bar, 50 μm.

**S4 Fig.**
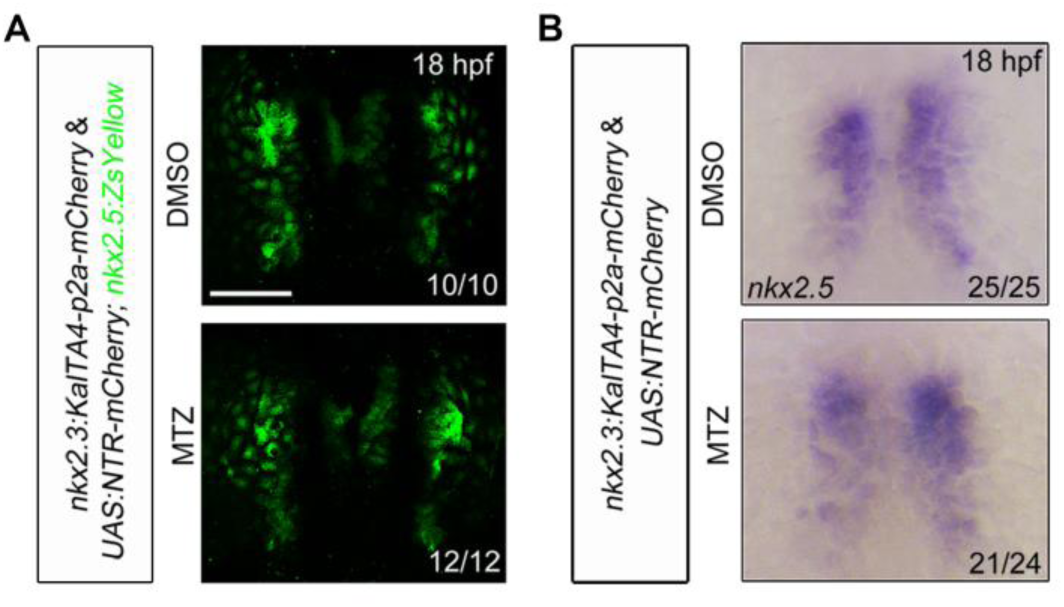
Pouch endoderm is not required for the segregation of *nxk2.5^+^* pharyngeal lineage from cardiac precursors. (A) Dorsal view of *Tg(nkx2.5:ZsYellow& nkx2.3:KalTA4-p2a-mCherry;UAS:NTR-mCherry)* embryos at 18 hpf, which were treated with DMSO or 10 mM MTZ from bud stage. Scale bar, 100 μm. (B) The *nkx2.5* transcripts were evaluated by *in situ* hybridizations in *Tg(nkx2.3:KalTA4-p2a-mCherry;UAS:NTR-mCherry)* embryos at 18 hpf, which were treated with DMSO or 10 mM MTZ from bud stage.

**S5 Fig.**
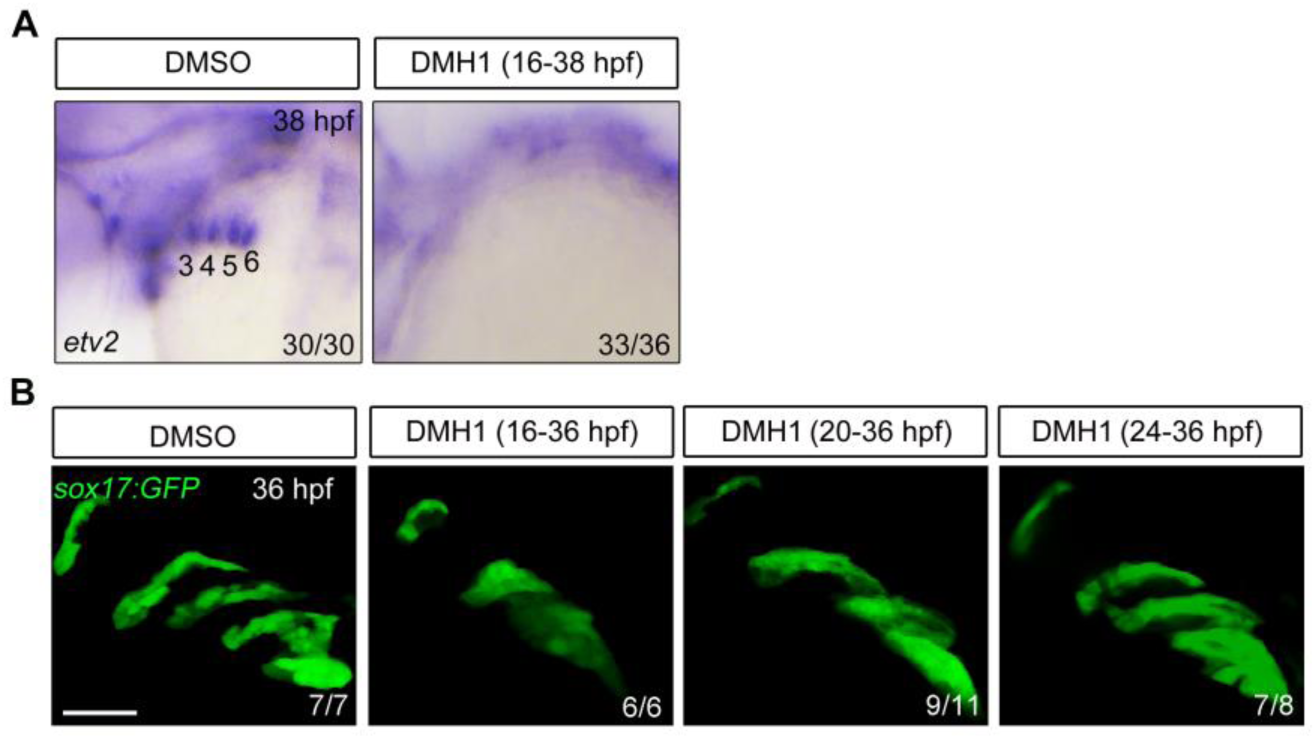
The effects of DMH1 treatment on PAA angioblast formation and pharyngeal pouch morphogenesis. (A) Expression analysis of PAA angioblast marker *etv2* by *in situ* hybridization at 38 hpf. The embryos were treated with DMSO or 10 μM DMH1 from 16 hpf. Numbers indicate the PAA angioblast clusters. (B) Lateral view of live *Tg(sox17:GFP)* embryos in different treatment groups at 36 hpf. Scale bar, 50 μm.

**S6 Fig.**
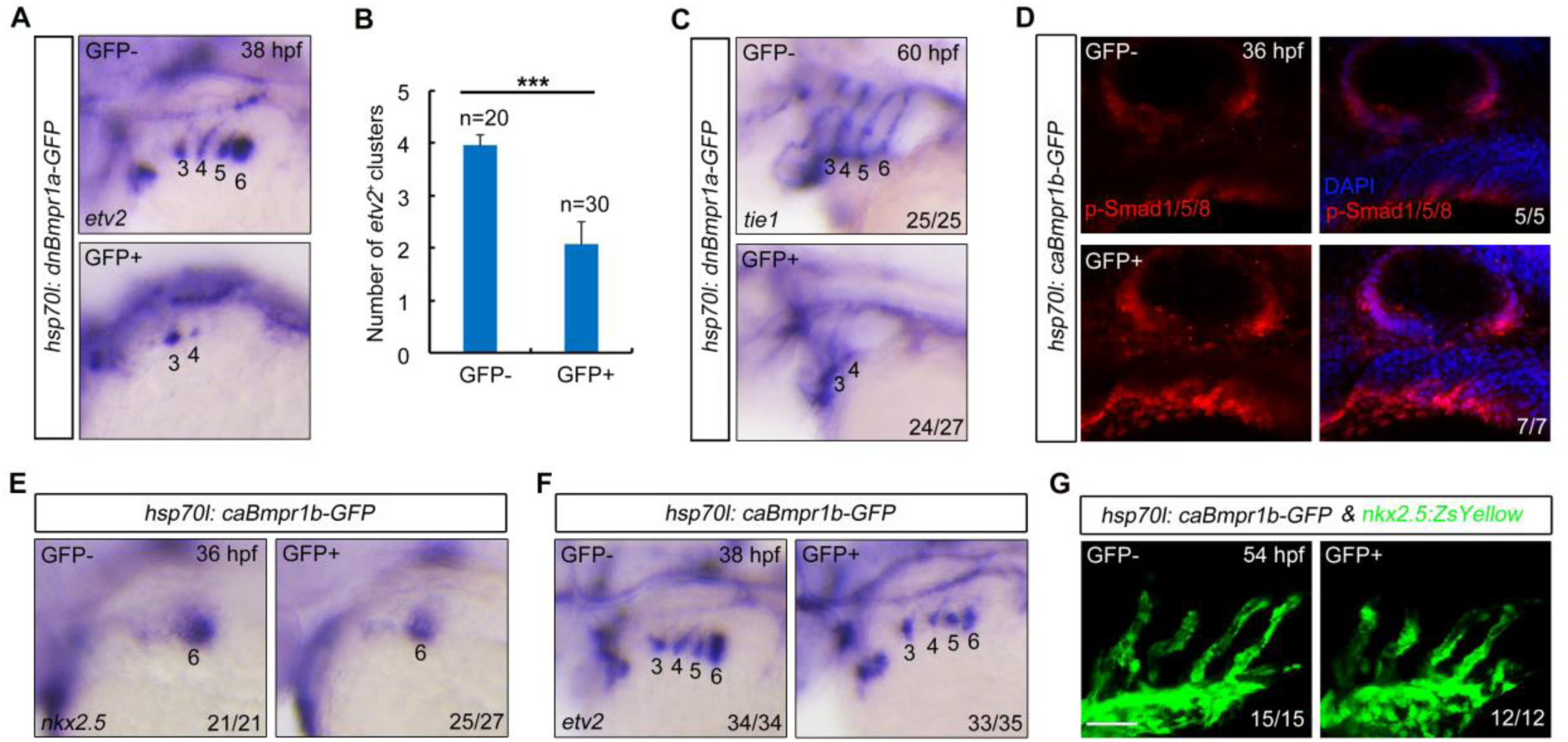
BMP signaling is necessary but not sufficient for PAA progenitor specification. (A-C) Expression analysis of *etv2* and *tie* by *in situ* hybridizations in *Tg(hsp70l:dnBmpr1a-GFP)* heated embryos. The embryos were heat shocked at 24 hpf for 20 min, and then harvested at the indicated developmental stages for *in situ* hybridizations. Numbers indicate the PAA angioblast clusters or PAAs. The average numbers of *etv2^+^* PAA angioblast clusters were quantified from three independent experiments and presented in (C). Student’s *t*-test, \*\*\**p* < 0.001. (D-F) Overactivation of BMP signal has no effect on PAA progenitor specification and angioblast differentiation. *Tg(hsp70l:caBmpr1b-GFP)* embryos were heat shocked at 24 hpf for 20 min, and then harvested at the indicated developmental stages for immunostaining (D) or *in situ* hybridizations (E and F). Scale bar, 50 μm. (G) Immunostaining of ZsYellow in *Tg(nkx2.5:ZsYellow;hsp70l:caBmpr1b-GFP)* embryos at 54 hpf, which were heat shocked at 24 hpf for 20 min. Scale bar, 50 μm.

**S7 Fig.**
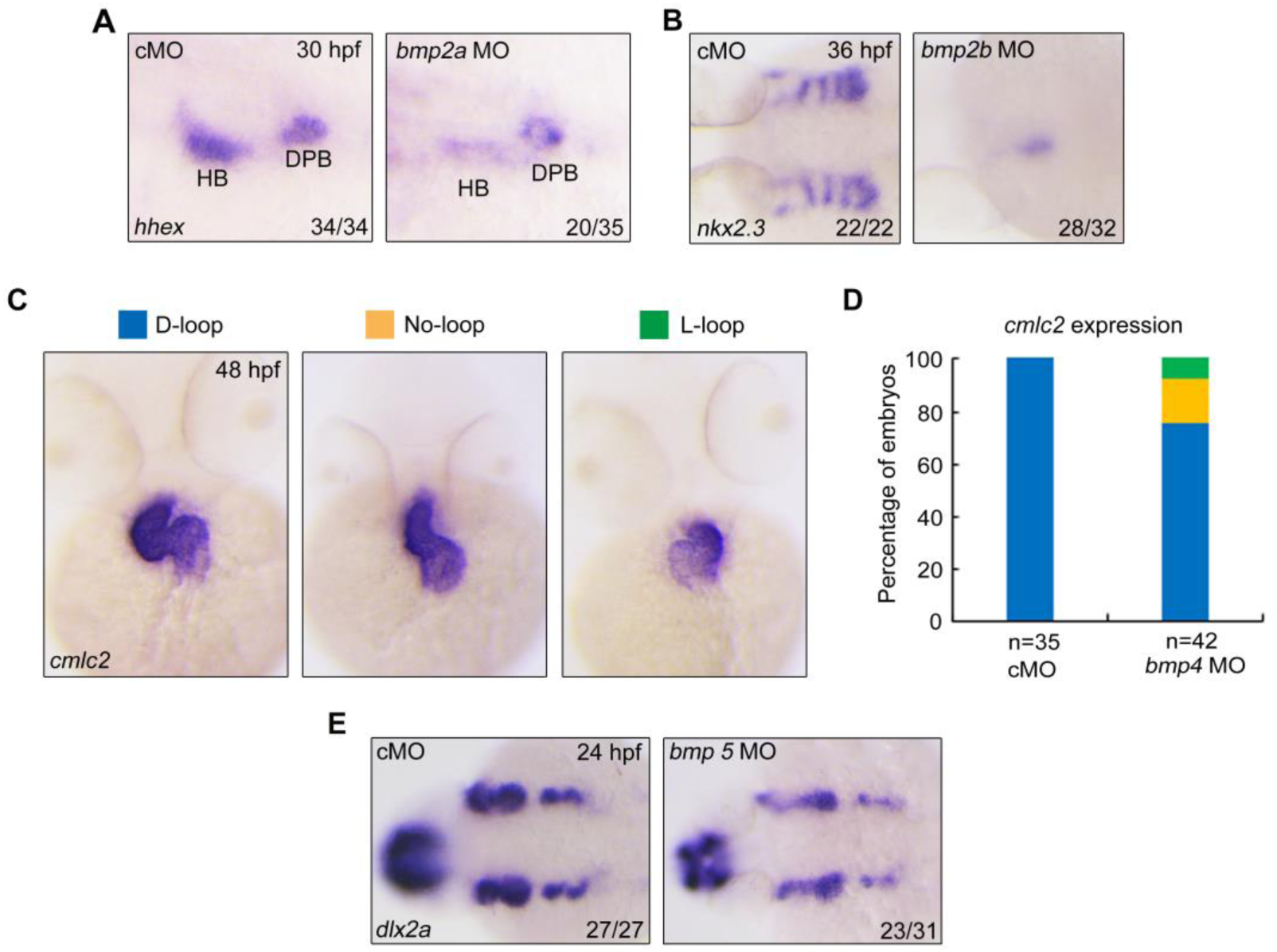
Validation the effects of the MOs targeting *bmp2a*, *bmp2b*, *bmp4*, and *bmp5* on embryonic development. (A) The expression of *hhex* was clearly reduced in the hepatic bud but not in the dorsal pancreatic bud in embryos injected with 2 ng *bmp2a* MO. HB, hepatic bud. DPB, dorsal pancreatic bud. (B) *In situ* hybridization of *nkx2.3* embryos injected with 0.3 ng *bmp2b* MO. Note that the expression of *nkx2.3* was nearly abolished in *bmp2b* morphants. (C-D) Knockdown of *bmp4* led to heart-looping defects. The embryos were injected with 2 ng *bmp4* MO at the one-cell stage, then were harvested at 48 hpf for *in situ* hybridization with *cmlc2* probe. Embryos with different phenotypes were shown in (C), and the ratios of the phenotypes were shown in (D). (E) Analysis of the expression of cranial neural crest cell marker *dlx2a* in embryos injected with 4 ng *bmp5* MO by *in situ* hybridization.

**S8 Fig.**
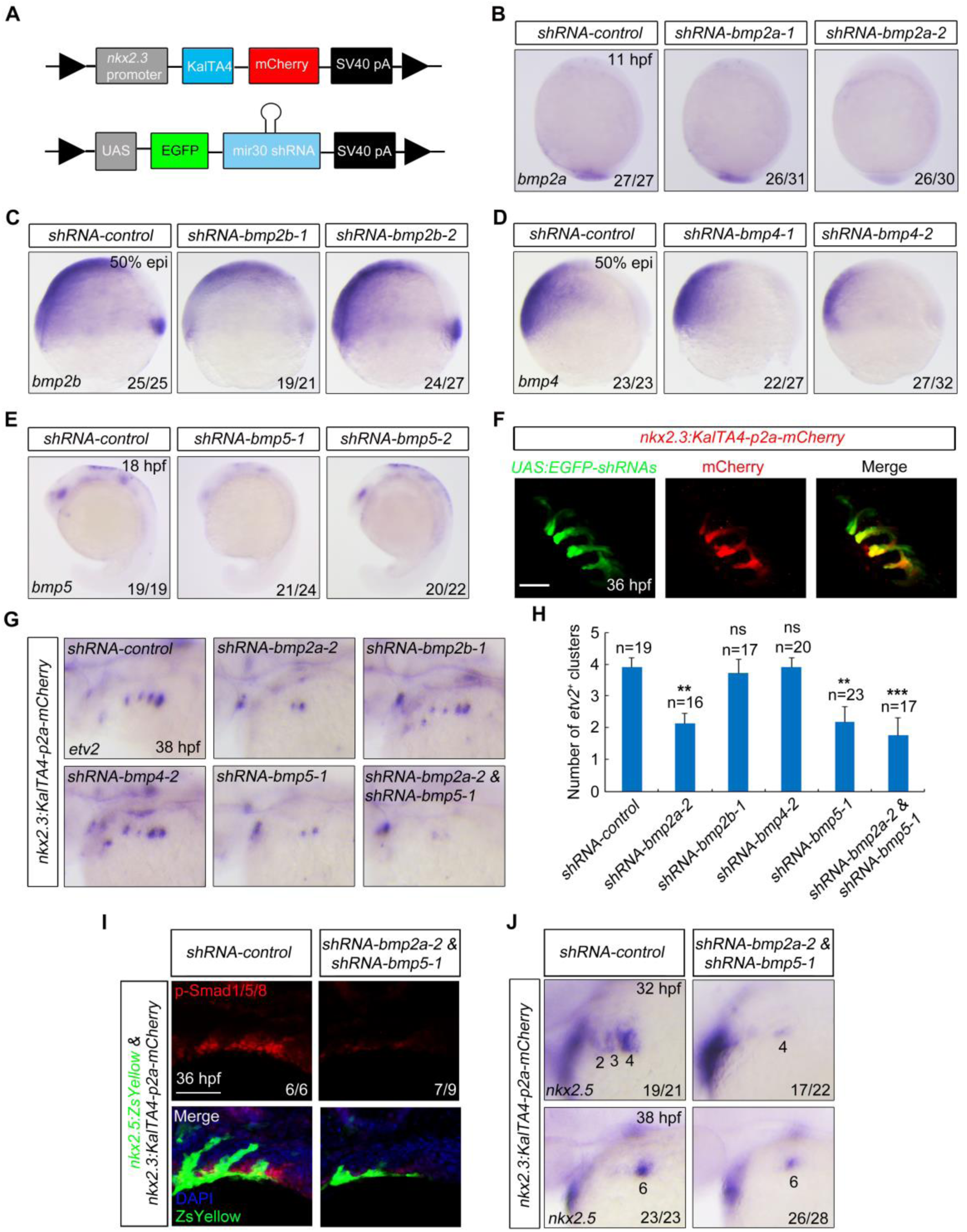
BMP2a and BMP5 function redundantly in PAA progenitor specification. (A) Schematic representation of the *nkx2.3:KalTA4-p2a-mCherry* and *UAS:miR30-shRNA* plasmid structures. (B-E) The effectiveness of the miR30-based shRNAs. Wild-type embryos were injected with 70 pg *KalTA4-p2a-mCherry* mRNA and 30 pg indicated *UAS:miR30-shRNA* plasmids at one-cell stage, and then subjected to *in situ* hybridizations for *bmp2a* expression at 11 hpf (B), for *bmp2b* and *bmp4* expression at 50% epiboly (50% epi) stage (C, D), and for *bmp5* expression at 18 hpf (E). (F) Live confocal images of *Tg(nkx2.3:KalTA4-p2a-mCherry)* embryos injected with 30 pg *UAS:EGFP-shRNA* plasmids and 200 pg Tol2 transposase mRNA at the one-cell stage. Scale bar, 50 μm. (G-H) *In situ* hybridization analysis of *etv2* expression in *Tg(nkx2.3:KalTA4-p2a-mCherry)* embryos at 38 hpf, which were injected with 200 pg Tol2 transposase mRNA and 30 pg indicated *UAS:EGFP-shRNA* plasmids at the one-cell stage (G). The average numbers of *etv2^+^* PAA angioblast clusters were quantified from three independent experiments and the group values were expressed as mean ± SD (H). Student’s *t*-test, \**p* < 0.05; \*\**p* < 0.01; \*\*\**p* < 0.001; ns, no significant difference. (I-J) Knockdown the expression of *bmp2a* and *bmp5* disrupts BMP signal activation in the pharyngeal mesoderm and the PAA progenitor specialization. *Tg(nkx2.5:ZsYellow;nkx2.3:KalTA4-p2a-mCherry)* embryos were injected with 200 pg Tol2 transposase mRNA and 30 pg indicated *UAS:EGFP-shRNA* plasmids at the one-cell stage. The resulting embryos were harvested at indicated developmental stages for immunostaining (I) or *in situ* hybridization (J) analysis. Scale bar, 50 μm.

**S9 Fig.**
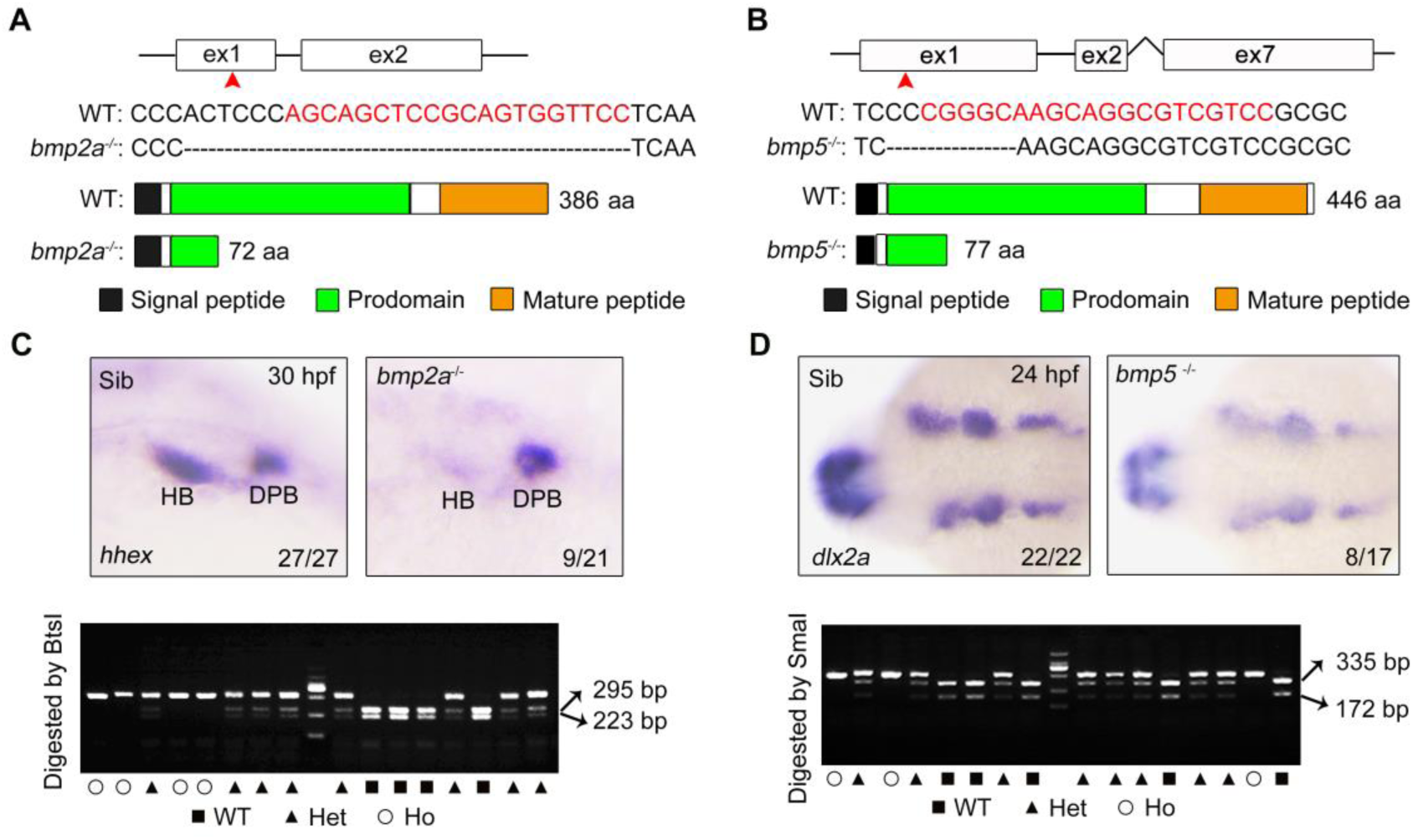
Generation of *bmp2a* and *bmp5* mutants. (A-B) Schematic diagram of *bmp2a*^-/-^ (A) and *bmp5*^-/-^ (B). The gRNA targeted exon 1 of *bmp2a*, causing a 26-base pair deletion, resulting in a premature stop codon that formed a truncated protein with 72 amino acids. The gRNA targeted exon 1 of *bmp5*, causing a 7-base deletion, and resulting in a premature stop codon that formed a truncated protein with 77 amino acids. The red asterisk indicates the relative position of the gRNA target sites and the red base indicates the gRNA target sequences. (C-D) *bmp2a*^-/-^ and *bmp5*^-/-^ mutants exhibited developmental defects in the formation of hepatic bud and cranial neural crest cell, respectively. Embryos generated by inter-crossing heterozygous *bmp2a* (C) or *bmp5* (D) mutant adult fishes were subjected to *in situ* hybridization (the upper panels), and then PCR genotyping analyses were conducted in the resulting embryos (the lower panels). Failure to digest with the indicated restriction enzymes is indicative of a possible target site deletion.

**S10 Fig.**
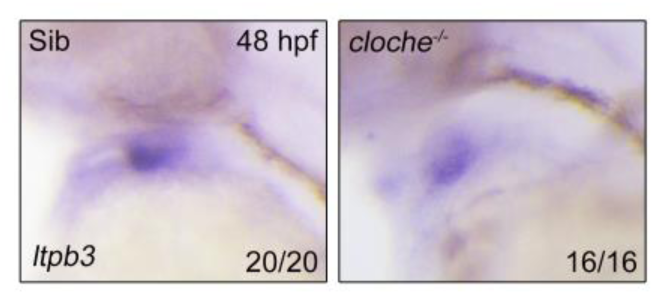
The development of the heart outflow tract is not impaired in *cloche* mutant. The expression analysis of *ltpb3* in *cloche* mutants and their siblings at 48 hpf by *in situ* hybridization.

**S1 Table.**
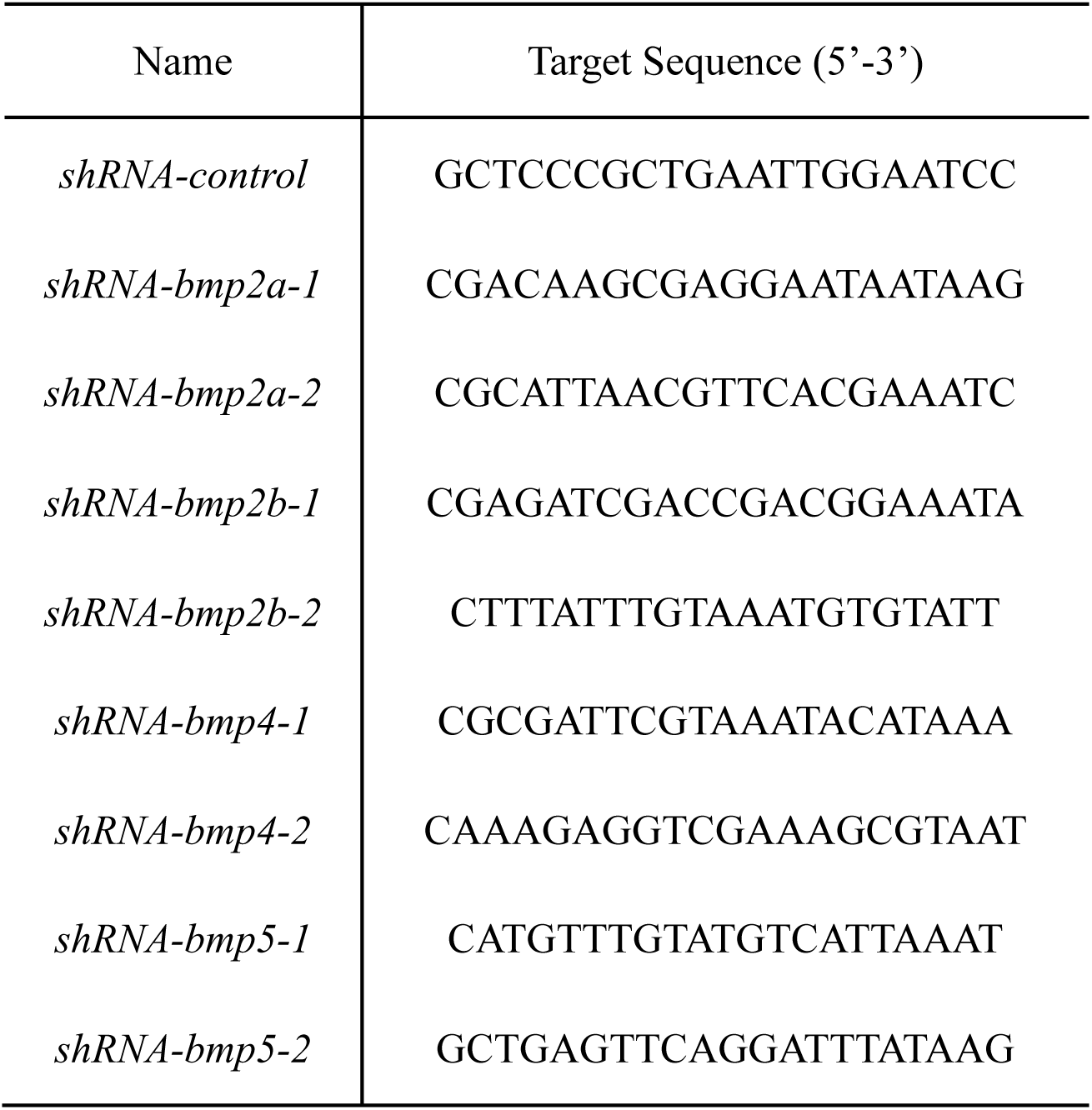
The target sequences of miR30-based shRNAs.

